# Observation weights to unlock bulk RNA-seq tools for zero inflation and single-cell applications

**DOI:** 10.1101/250126

**Authors:** Koen Van den Berge, Fanny Perraudeau, Charlotte Soneson, Michael I. Love, Davide Risso, Jean-Philippe Vert, Mark D. Robinson, Sandrine Dudoit, Lieven Clement

## Abstract

Dropout events in single-cell transcriptome sequencing (scRNA-seq) cause many transcripts to go undetected and induce an excess of zero read counts, leading to power issues in differential expression (DE) analysis. This has triggered the development of bespoke scRNA-seq DE methods to cope with zero inflation. Recent evaluations, however, have shown that dedicated scRNA-seq tools provide no advantage compared to traditional bulk RNA-seq tools. We introduce a weighting strategy, based on a zero-inflated negative binomial (ZINB) model, that identifies excess zero counts and generates gene and cell-specific weights to unlock bulk RNA-seq DE pipelines for zero-inflated data, boosting performance for scRNA-seq.

## Background

Transcriptomics has become one of the standard tools in modern biology to unravel the molecular basis of biological processes and diseases. One of the most common applications of transcriptome profiling is the discovery of *differentially expressed* (DE) genes, which exhibit changes in expression levels across conditions [Love et al., 2014, Robinson et al., 2010, Law et al., 2014]. Over the last decade, transcriptome sequencing (RNA-seq) has become the standard technology for transcriptome profiling, enabling researchers to study average gene expression over bulks of thousands of cells [Wang et al., 2009, Goodwin et al., 2016]. The advent of single-cell RNA-seq (scRNA-seq) enables high-throughput transcriptome profiling at the resolution of single cells and allows, among other things, research on cell developmental trajectories, cell-to-cell heterogeneity, and the discovery of novel cell types [Lonnberg et al., 2017, Buettner et al., 2015, Patel et al., 2014, Kolodziejczyk et al., 2015a, Li et al., 2017, Usoskin et al., 2014].

In scRNA-seq, individual cells are first captured, their RNA is then reverse-transcribed into cDNA, which is greatly amplified from the minute amount of starting material, and the resulting library is finally sequenced [Kolodziejczyk et al., 2015b]. Transcript abundances are typically estimated by counts, which represent the number of sequencing reads mapping to an exon, transcript, or gene. Many scRNA-seq protocols have been published to conduct such core steps [Nakamura et al., 2015, Wu et al., 2013, Islam et al., 2013, 2011, Picelli et al., 2014, Hashimshony et al., 2016], but despite these advances, scRNA-seq data remain inherently noisy. *Dropout* events cause many transcripts to go undetected for technical reasons, such as inefficient cDNA polymerization, amplification bias, or low sequencing depth, leading to an excess of zero read counts as compared to bulk RNA-seq data [Hashimshony et al., 2016, Finak et al., 2015]. In addition, excess zeros can also occur for biological reasons, such as transcriptional bursting Raj and van Oudenaarden, 2008]. There are therefore two types of zeros in scRNA-seq data: *biological zeros*, when a gene is simply not expressed in the cell, and *technical zeros* (i.e., dropouts), when a gene is expressed in the cell but not detected. *Zero inflation*, i.e., excess zeros compared to standard count distributions (e.g., negative binomial) used in bulk RNA-seq, occurs for both biological and technical reasons and disentangling the two sources is non-trivial. In addition, scRNA-seq counts are inherently more variable than bulk RNA-seq counts because the transcriptional signal is not averaged across thousands of individual cells (Additional File 1: Figure S1), making cell-to-cell heterogeneity, cell type mixtures, and stochastic expression bursts important contributors to between-sample variability [Raj et al., 2006, Buettner et al., 2015].

Typical scRNA-seq data analysis workflows often involve identifying cell types *in silico* using tailored clustering algorithms [Pierson and Yau, 2015, Risso et al., 2017] or ordering cells along developmental trajectories, where cell types are defined as terminal states of the developmental process [Setty et al., 2016, Qiu et al., 2017, Lonnberg et al., 2017, Street et al., 2017]. A natural subsequent step is the discovery of marker genes for the defined cell types by assessing differential gene expression between these groups. Another common setting is the identification of marker genes for *a priori* known cell types. Differential expression analysis between homogeneous cell populations as in the aforementioned scRNA-seq applications is the use case for our method.

Popular bulk RNA-seq DE tools, such as those implemented in the Bioconductor R packages edgeR [Robinson et al., 2010] and DESeq2 [Love et al., 2014], assume a negative binomial (NB) count distribution across biological replicates, while limma-voom [Law et al., 2014] uses linear models for log-transformed counts and observation-level weights to account for the mean-variance relationship of the transformed count data. Such tools can also be applied for scRNA-seq DE analysis [Lun et al., 2016]. However, dropouts, transcriptional bursting, and high variability in scRNA-seq data raise concerns about their validity. This has triggered the development of novel dedicated tools, which typically introduce an additional model component to account for the excess of zeros through, for example, zero-inflated (`SCDE`, Kharchenko et al. [2014]) or hurdle (`MAST`, Finak et al. [2015]) models. However, Jaakkola et al. [2016] and Soneson and Robinson [2017a] have recently shown that these bespoke tools do not provide systematic benefits over standard bulk RNA-seq tools in scRNA-seq applications.

We argue that standard bulk RNA-seq tools, however, still suffer in performance due to zero inflation with respect to the negative binomial distribution. We illustrate this using biological coefficient of variation (BCV) plots [McCarthy et al., 2012a], which represent the mean-variance relationship of the counts. Note that the BCV plots of scRNA-seq data exhibit striped patterns (Figure 1a-b and Additional File 1: Figure S2 for scRNA-seq datasets subsampled to ten cells) that are indicative of genes with few positive counts (Additional File 1: Figure S3) and very high dispersion estimates. Randomly adding zeros to bulk RNA-seq data, likewise consisting of ten samples, also results in similar striped patterns (Figure 1c-d). Negative binomial models, as implemented in `DESeq2` and `edgeR`, will thus accommodate excess zeros by overestimating the dispersion parameter, which jeopardizes the power to infer differential expression. However, by correctly identifying the excess zeros and downweighting them in the dispersion estimation and model-fitting, one reconstructs the original mean-variance relationship (Figure 1e), thus recovering the power to detect differential expression (Figure 1f). Hence, identifying and downweighting excess zeros provides the key to unlocking bulk RNA-seq tools for scRNA-seq differential expression analysis. Note that methods based on a zero-inflated negative binomial (ZINB) model naturally implement such an approach: Excess zeros are attributed weights through the zero inflation probability and inference can be focused on the mean of the negative binomial count component.

**Figure 1:**
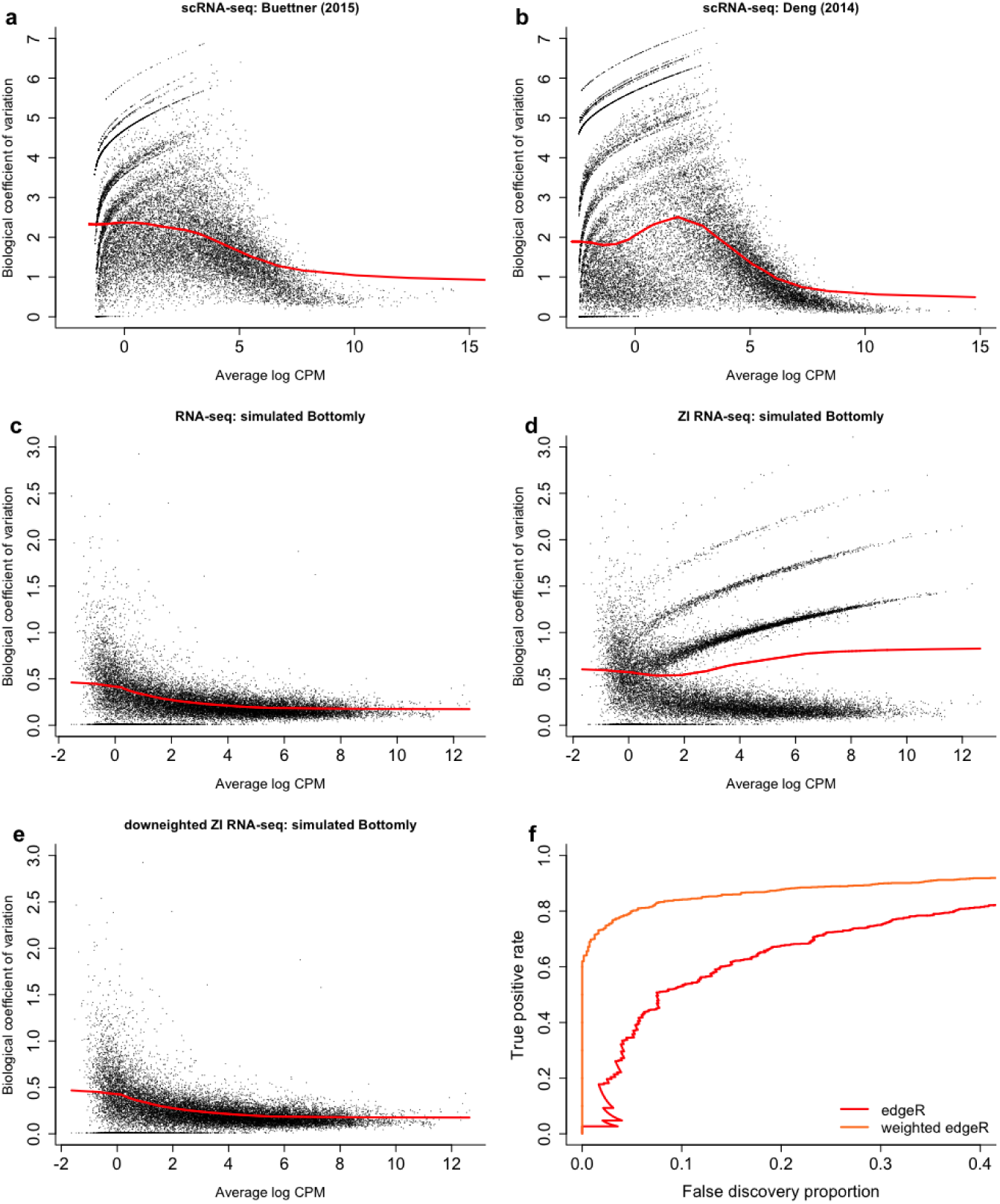
Zero inflation results in overestimated dispersion and jeopardizes power to discover differentially expressed genes. (a–e) Scatterplots of estimated biological coefficient of variation (BCV, defined as the square root of the negative binomial dispersion parameter *φ*) against average log count per million (CPM) computed using `edgeR`. (a) BCV plot for the real Buettner et al. [2015] scRNA-seq dataset subsampled to *n* = 10 cells. (b) BCV plot for the real Deng et al. [2014] scRNA-seq dataset subsampled to *n* = 10 cells. Both panels (a) and (b) show striped patterns in the BCV plot, which significantly distort the mean-variance relationship, as represented by the red curve. (c) BCV plot for a simulated bulk RNA-seq dataset (*n* = 10), obtained from the Bottomly et al. [2011] dataset using the simulation framework of Zhou et al. [2014]. Dispersion estimates generally decrease smoothly as gene expression increases. (d) BCV plot for a simulated zero-inflated bulk RNA-seq dataset, obtained by randomly introducing 5% excess zero counts in the dataset from (c). Zero inflation leads to overestimated dispersion for the genes with excess zeros, resulting in striped patterns, as observed also for the real scRNA-seq data in panels (a) and (b). (e) BCV plot for simulated zero-inflated bulk RNA-seq dataset from (d), where excess zeros are downweighted in dispersion estimation (i.e., weights of 0 for excess zeros and 1 otherwise). Downweighting recovers the original mean-variance trend. (f) True positive rate vs. false discovery proportion (FDP-TPR curves) for simulated zero-inflated dataset of (d). The performance of `edgeR` (red curve) is deteriorated in a zero-inflated setting due to overestimation of the dispersion parameter. However, assigning the excess zeros a weight of zero in the dispersion estimation and model-fitting results in a dramatic performance boost (orange curve). Hence, downweighting excess zero counts is the key to unlocking bulk RNA-seq tools for zero inflation.

We therefore propose a weighting strategy based on ZINB models to unlock bulk RNA-seq tools for scRNA-seq DE analysis. In this manuscript, we build on the zero-inflated negative binomial-based wanted variation extraction (ZINB-WaVE) method of Risso et al. [2017], designed specifically for scRNA-seq data. ZINB-WaVE efficiently identifies excess zeros and provides gene and cell-specific weights to unlock bulk RNA-seq pipelines for zero-inflated data. As most bulk RNA-seq DE methods are based on generalized linear models (GLM), which readily accommodate observation-level weights, our approach seamlessly integrates with standard pipelines (e.g., `edgeR`, `DESeq2`, `limma`). Our method is shown to outperform competing methods on simulated bulk and single-cell RNA-seq datasets. We also illustrate our method on two publicly available real datasets. As detailed in the “Software implementation” section, our approach is implemented in open-source Bioconductor R packages and the code for reproducing the analyses presented in this manuscript is provided in a GitHub repository.

## Results

### ZINB-WaVE extends bulk RNA-seq tools to handle zero-inflated data

We argue that standard bulk RNA-seq methods for inferring differential gene expression suffer from zero inflation with respect to the assumed negative binomial distribution when applied to scRNA-seq data. We propose instead to model scRNA-seq data using a zero-inflated model and perform inference on the count component of the model, which is equivalent to standard NB regression where excess zeros are downweighted based on posterior probabilities (weights) inferred from a ZINB model. Such weights play a central role in many estimation approaches for ZINB models (e.g., [A. and Trivedi, 2013]). In this contribution, we show that the weights can effectively unlock bulk RNA-seq methods for zero-inflated data, allowing us, in particular, to borrow strength across genes to estimate dispersion parameters. Here, we use weights derived from the zeroinflated negative binomial-based wanted variation extraction (ZINB-WaVE) method of Risso et al. [2017], which is a general and flexible framework for the extraction of low-dimensional signal from scRNA-seq read counts, accounting for zero inflation (i.e., dropouts, bursting), over-dispersion, and the discrete nature of the data. Note that although we focus on ZINB-WaVE weights, our weighted DE approach is generic and researchers might choose to adopt their own weights.

A zero-inflated negative binomial (ZINB) distribution is a two-component mixture between a point mass at zero and a negative binomial distribution. Specifically, the density function for the ZINB-WaVE model is

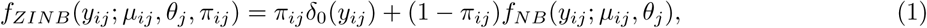

where *y*_*ij*_ denotes the read count for cell *i* and gene *j*, *π*_*ij*_ the mixture probability for zero inflation, *f*_*NB*_(·; *μ*_*ij*_, *θ*_*j*_) the negative binomial probability mass function with mean *μ*_*ij*_ and dispersion *θ*_*j*_, and *δ*_0_ the Dirac delta function (see Equations (3) and (4)).

The ZINB-WaVE parameterization of the NB mean *μ* and ZI probability *π* in Equation (4) allows adjusting for both known (e.g., treatment, batch, quality control measures) and unknown (RUV) [Gagnon-Bartsch and Speed, 2012, Risso et al., 2014] cell-level covariates, i.e., supervised and unsupervised normalization, respectively. It also allows adjusting for known gene-level covariates (e.g., length, GC-content). The ZINB-WaVE model and its associated penalized maximum likelihood estimation procedure are described more fully in the “Methods” section and in Risso et al. [2017].

From the ZINB-WaVE density of Equation (1), one can readily derive the posterior probability that a count *y*_*ij*_ was generated from the negative binomial count component

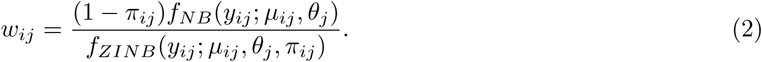

We propose to use these probabilities as weights in bulk RNA-seq DE analysis methods, such as those implemented in the Bioconductor R packages `edgeR`, `DESeq2`, and limma (limma-voom method with voom function). All of these methods are based on the methodology of generalized linear models, which readily accommodates inference based on observation-level weights. Note that although the ZINB-WaVE weights are gene and cell-specific, the GLMs are fit gene by gene; hence, for a given gene, the cell-specific weights are used as observation-specific weights in the GLMs. The implementation of the weighting strategy for `edgeR`, `DESeq2`, and limma-voom is described in greater detail in the “Methods” section.

### Impact of zero inflation on mean-variance relationship

We have already noted that adding zeros to bulk RNA-seq data results in an overestimation of the dispersion parameter. This leads to striped patterns in the BCV plot (Figure 2a), which are indicative of genes with many zeros (Additional File 1: Figure S3) and very high dispersion estimates. Our ZINB-WaVE method, however, identifies many of the introduced excess zeros as such (Figure 2a-b), by classifying them in the zero-inflated component of the ZINB mixture distribution. Using our posterior probabilities as observation-level weights in `edgeR` recovers the original BCV plot and mean-variance trend (Figure 2c), illustrating the ability of our method to account for zero inflation. Hence, observation weights provide the key to unlocking standard bulk RNA-seq tools for zero-inflated data.

**Figure 2:**
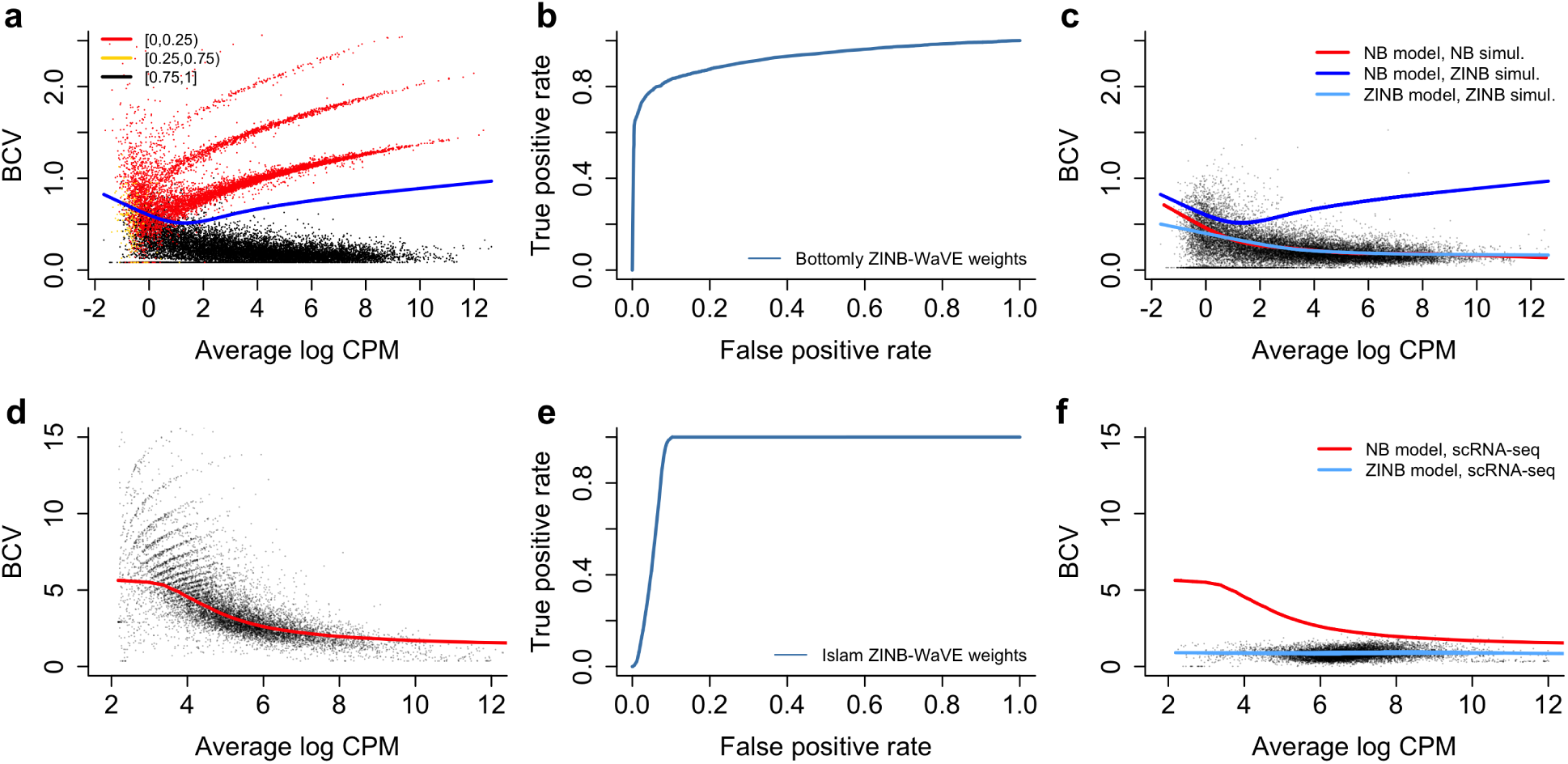
Impact of zero inflation on mean-variance relationship for simulated bulk RNA-seq and Islam scRNA-seq datasets. Zero inflation distorts the mean-variance trend in (sc)RNA-seq data, but is correctly identified by the ZINB-WaVE method. The top panels represent simulated data based on the Bottomly et al. [2011] bulk RNA-seq dataset (as in Figure 1), for a two-group comparison with five samples in each group, where 5% of the counts were randomly replaced by zeros. The bottom panels represent the scRNA-seq dataset from Islam et al. [2011]. (a) The BCV plot shows that randomly replacing 5% of the read counts with zeros induces zero inflation and distorts the mean-variance trend by leading to overestimated dispersion parameters. Points are color-coded according to the average ZINB-WaVE posterior probability for all zeros for a given gene and the blue line represents the mean-variance trend estimated with `edgeR`. (b) Receiver operating characteristic (ROC) curve for the identification of excess zeros by the ZINB-WaVE method. A very good classification precision is obtained. (c) Downweighting excess zeros using the ZINB-WaVE posterior probabilities recovers the original mean-variance trend (as indicated with the red line) and inference on the negative binomial count component will now no longer be biased because of zero inflation. The light blue line represents the estimated mean-variance trend for ZINB-WaVE-weighted `edgeR`. The blue line is the trend estimated by unweighted `edgeR` on zero-inflated data as in panel (a). (d) The BCV plot for the Islam et al. [2011] dataset illustrates the higher variability of scRNA-seq data as compared to bulk RNA-seq data (note the difference in y-axis scales between (a) and (d)). As in (a), zero inflation induces striped patterns leading to an overestimation of the NB dispersion parameter. (e) ROC curve for the identification of excess zeros by the ZINB-WaVE method for scRNA-seq data simulated from the Islam dataset using the simulation framework described in “Methods”. A good classification precision is obtained, but note the difference with bulk RNA-seq data: The noisier scRNA-seq dataset makes excess zero identification harder. (f) Using the ZINB-WaVE posterior probabilities as observation weights results in lower estimates of the dispersion parameter, unlocking powerful differential expression analysis with standard bulk RNA-seq DE methods. Note that since many zeros are identified as excess, the scale of the BCV plot is now similar to that of a standard bulk RNA-seq dataset. The red line is the mean-variance trend for unweighted `edgeR`, as in panel (d), and the light blue line is the mean-variance trend for ZINB-WaVE-weighted `edgeR`. A similar pattern is observed for the simulated Islam dataset (Additional File 1: Figure S26).

The BCV plot for the Islam et al. [2011] scRNA-seq dataset (Figure 2d) shows similar striped patterns as for zero-inflated bulk RNA-seq data. Such patterns are observed in many single-cell datasets (Additional File 1: Figure S2). ZINB-WaVE identifies many zeros to be excess for the Islam dataset. It also provides good classification power for excess zeros for data simulated from the Islam dataset (Figure 2e). Incorporating the ZINB-WaVE weights in an `edgeR` analysis removes the striped patterns and yields a BCV plot that is similar to that for bulk RNA-seq data (Figure 2f), suggesting that zero inflation was indeed present and accounted for.

### High power and false positive control on simulated (sc)RNA-seq data

We provide a scRNA-seq data simulation paradigm that retains gene-specific characteristics as well as global associations across all genes (see “Methods” for details). More specifically, we first estimate dataset-specific associations between zero abundance, sequencing depth, and average log counts per million (CPM), and next explicitly account for these associations in our simulation model (Additional File 1: Figures S4–S5).

The scRNA-seq simulation study is based on three datasets: the Islam et al. [2011] dataset, comparing 48 embryonic stem cells to 44 embryonic fibroblasts in the mouse; a subset of the Trapnell et al. [2013] dataset, comparing differentiating human myoblasts at the 48h (85 cells) and 72h (64 cells) timepoints; and a 10x Genomics peripheral blood mononuclear cells (PBMC) dataset (see “Real datasets” section in “Methods” for details). The datasets differ in throughput, sequencing depth, and extent of zero inflation, e.g., Additional File 1: Figure S6 shows a higher proportion of excess zeros in the Islam dataset as compared to the Trapnell dataset, an observation further supported by the fact that the Islam and Trapnell datasets contain ~ 65% and ~ 48% zeros, respectively. 10x Genomics datasets are known to contain even more zeros; the evaluated subset of the PBMC dataset contains ~ 87% zeros. The simulated datasets successfully mimic the characteristics of the original datasets, as evaluated with the R package countsimQC [Soneson and Robinson, 2017b] (Additional Files 2–4). This diverse range of datasets is therefore representative of scRNA-seq datasets that occur in practice and a suitable basis for method evaluation and comparison.

We evaluate method performance in terms of sensitivity and false positive control using false discovery proportion - true positive rate (FDP-TPR) curves. Figure 3 (Additional File 1: Figure S7) illustrates that many methods break down on the simulated Islam dataset due to a high degree of zero inflation. Surprisingly, even methods specifically developed to deal with excess zeros, like `SCDE` and metagenomeSeq, suffer from poor performances, with `MAST` being a notable exception. The `DESeq2` methods, however, are able to cope with the high degree of zero inflation. Note that, in general, it is a good strategy to disable the outlier imputation step in `DESeq2`, since it deteriorates performance on scRNA-seq data (Additional File 1: Figure S8). Seurat, limma-voom, and `SCDE` have very low sensitivity. The methods based on ZINB-WaVE weights dominate all competitors in terms of sensitivity and specificity, providing high power, good false discovery rate (FDR) control, and sensible p-value distributions (Additional File 1: Figure S9). Note that the remaining methods also suffer from poor FDR control.

**Figure 3:**
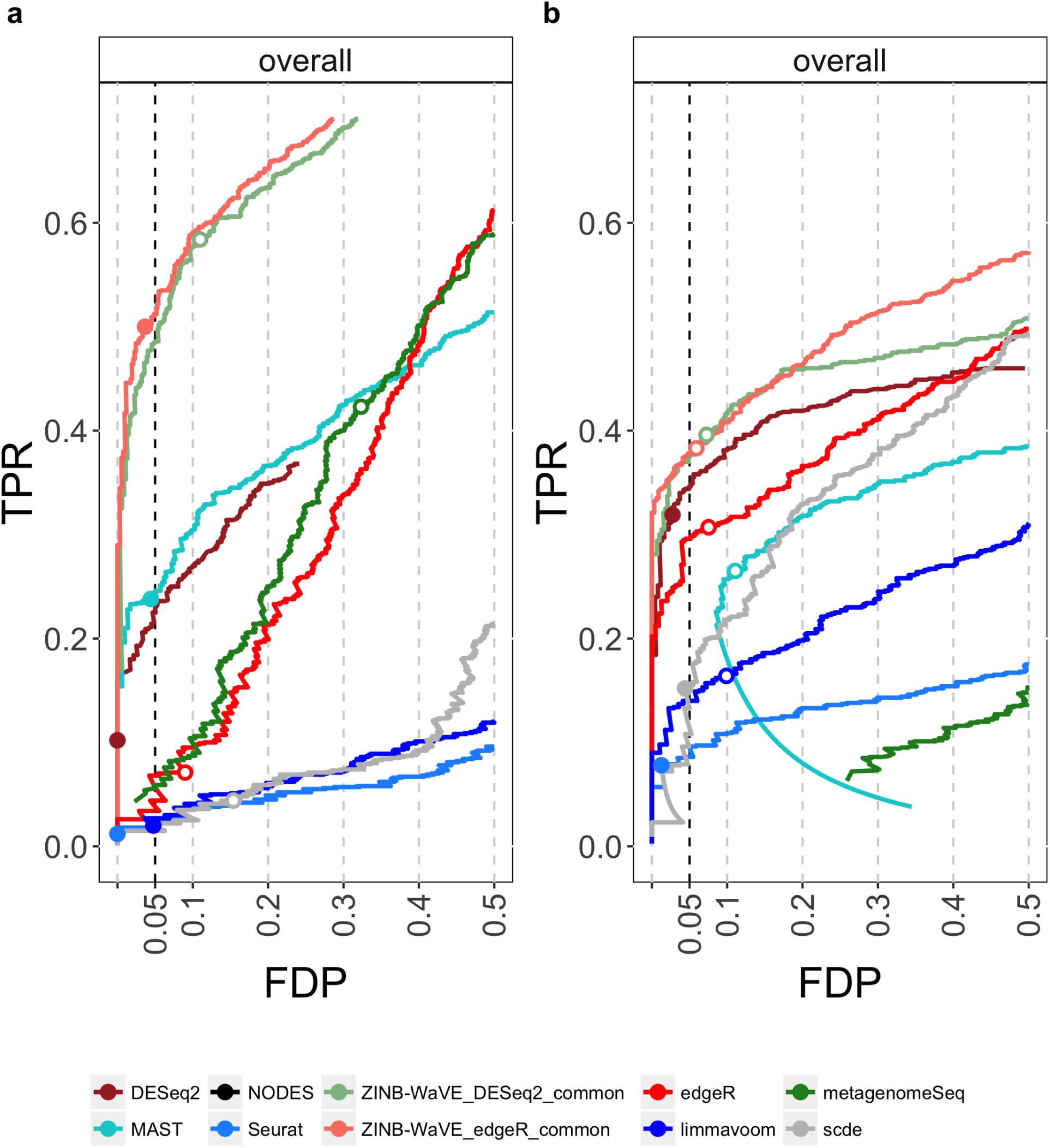
Comparison of DE methods on simulated scRNA-seq data. (a) scRNA-seq data simulated from Islam et al. [2011] dataset (*n* = 90). (b) scRNA-seq data simulated from Trapnell et al. [2013] dataset (*n* = 150). DE methods are compared based on scatterplots of the true positive rate (TPR) vs. the false discovery proportion (FDP); zoomed versions of the FDP-TPR curves are shown here, full curves are displayed in Additional File 1: Figure S7. Circles represent working points on a nominal 5% FDR level and are filled if the empirical FDR (i.e., FDP) is below the nominal FDR. Methods based on ZINB-WaVE weights clearly outperform other methods for both simulated datasets. Note that the methods differ in performance between datasets, possibly because of a higher degree of zero inflation in the Islam dataset. The `SCDE` and metagenomeSeq methods, specifically developed to deal with excess zeros, are outperformed in both simulations by ZINB-WaVE-based methods and by `DESeq2`. The `DESeq2` curve in panel (a) is cut off due to NA adjusted *p*-values resulting from independent filtering. The behavior in the lower half of the curve for `MAST` in (b) is due to a smooth increase in true positives with an identical number of false positives over a range of low FDR cut-offs. The curve for NODES is not visible on this figure, only in the full FDP-TPR curves.

Since zero inflation is fairly modest for the Trapnell dataset, most methods perform better than for the Islam simulation (Figure 3). The ZINB-WaVE-based methods and `DESeq2` outperform the remaining methods in terms of sensitivity and provide good FDR control. `edgeR` is their closest competitor and the remaining methods provide much lower sensitivity and/or very liberal FDR control. Note how bespoke scRNA-seq methods seem to break down on datasets with a lower degree of zero inflation, often providing too liberal or too conservative p-value distributions, while ZINB-WaVE-based methods in general show a reasonable p-value distribution, with an enrichment of low *p*-values and approximately uniformly distributed larger p-values (Additional File 1: Figure S10).

Typical 10x Genomics datasets contain a high number of cells with shallow sequencing depth, due to the extreme multiplexing of libraries. As a result, counts and hence estimated NB means are lower, making zeros more plausible according to the NB distribution and excess zeros thus harder to identify. This is picked up by the simulation framework, where only ~ 8% of the genes were simulated to have at least one excess zero in *n* = 1,200 samples. Bulk RNA-seq methods can hence be expected to be among the top performers. Figure 4 shows FDP-TPR curves for the 10x Genomics simulation study, demonstrating a good performance of bulk RNA-seq methods `edgeR` and `DESeq2`. ZINB-WaVE `edgeR` and ZINB-WaVE `DESeq2` are among the top performers, having comparable or slightly lower performance as compared to their unweighted counterparts. `MAST` is their closest competitor, providing good sensitivity and FDR control. `SCDE`, `NODES`, metagenomeSeq, and limma-voom have lower sensitivity and/or very liberal FDR control as compared to the dominating methods. These results suggest that, in a scenario of low counts or low degree of zero inflation, ZINB-WaVE-weighted `edgeR`/`DESeq2` reduce to standard unweighted `edgeR`/`DESeq2`, while other bespoke scRNA-seq tools may deteriorate in performance. This notion is further supported by results on simulated bulk RNA-seq data, where ZINB-WaVE-weighted `edgeR`/`DESeq2` have a similar performance as standard unweighted `edgeR`/`DESeq2` in the absence of zero inflation (Additional File 1: Figure S11). Hence, adopting ZINB-WaVE-based DE methods provides a performance boost in zero-inflated applications, while performance is similar in the absence of zero inflation.

**Figure 4:**
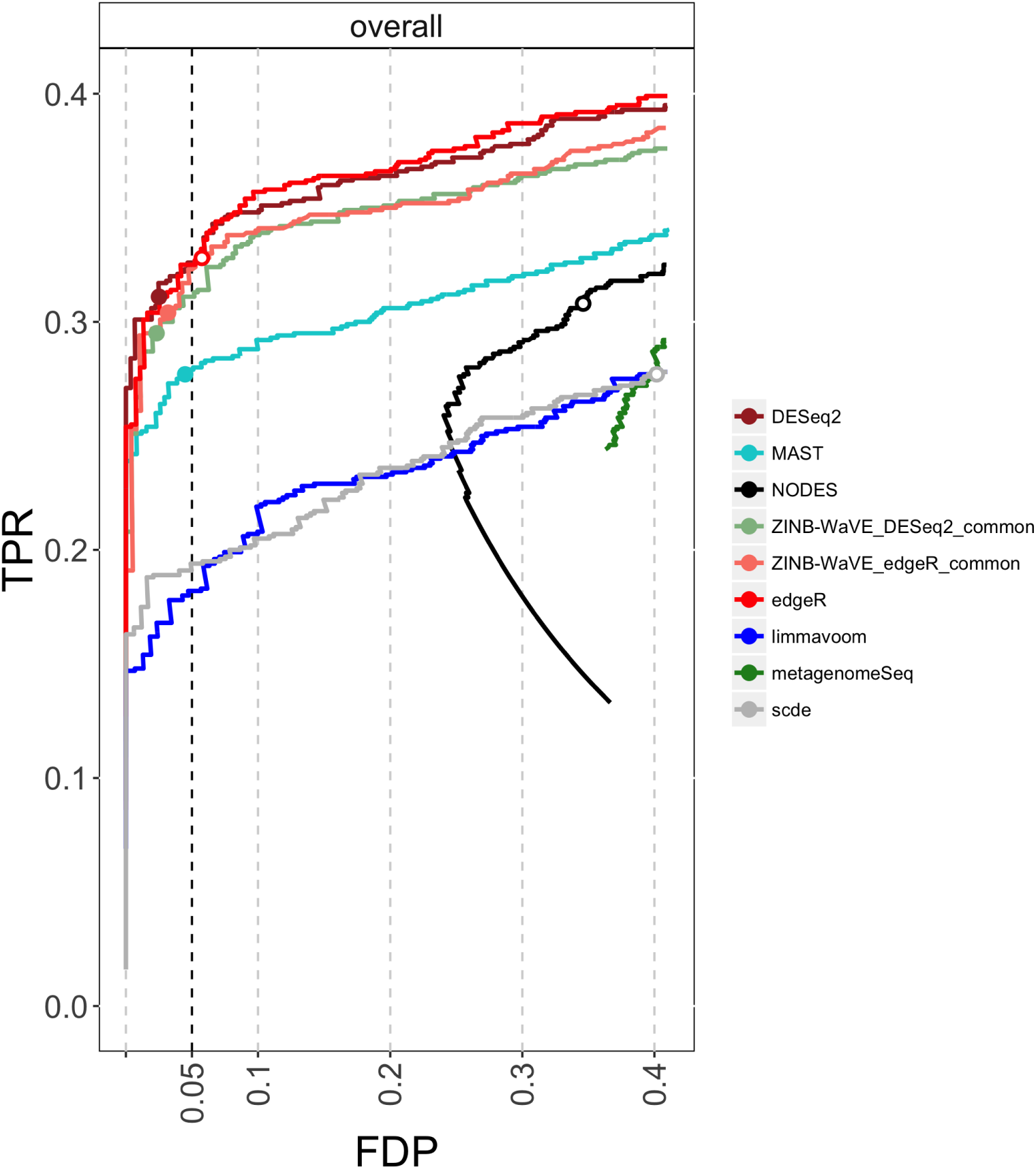
Comparison of DE methods on simulated scRNA-seq datasets. DE methods are compared based on FDP-TPR curves for data simulated from a 10x Genomics PBMC scRNA-seq dataset (*n* = 1, 200); zoomed versions of the FDP-TPR curves are shown here, full curves are displayed in Additional File 1: Figure S12. Circles represent working points on a nominal 5% FDR level and are filled if the empirical FDR (i.e., FDP) is below the nominal FDR. 10x Genomics sequencing typically involves high-throughput and massive multiplexing, resulting in very shallow sequencing depths and thus low counts, making it extremely difficult to identify excess zeros. Unweighted and ZINB-WaVE-weighted `edgeR` are tied for best performance, followed by ZINB-WaVE-weighted `DESeq2`. In general, bulk RNA-seq methods are performing well in this simulation, probably because the extremely high zero abundance in combination with low counts can be reasonably accommodated by the negative binomial distribution. The behavior in the lower half of the curve for NODES is due to a smooth increase in true positives with an identical number of false positives over a range of low FDR cut-offs.

All analyses performed in this work are based on estimating one common dispersion parameter across all genes for the ZINB-WaVE model. ZINB-WaVE allows the estimation of genewise dispersion parameters, however, this approach is much more computationally intensive and can be an order of magnitude slower. Additional File 1: Figures S7 and S12 show that estimating genewise dispersion parameters does not seem to be required for calculating the ZINB-WaVE weights, since no gain in performance is achieved when doing so. Note that genewise dispersions are still estimated by `edgeR` and `DESeq2` in the final DE inference procedure.

### False positive rate control

We compared our ZINB-WaVE-weight-based method to commonly-used DE methods for mock comparisons based on two publicly available real scRNA-seq datasets. We assessed performance based on the per-comparison error rate (PCER), defined as the proportion of false positives (i.e., Type *i* errors) among all genes being considered for DE, where a gene is declared DE if its nominal unadjusted p-value is less than or equal to 0.05.

The first dataset, referred to as Usoskin [Usoskin et al., 2014] dataset, concerns 622 mouse neuronal cells from the dorsal root ganglion, classified in eleven categories. The authors acknowledge the existence of a batch effect related to the picking session for the cells. We find that the batch effect is not only associated with expression measures, but also influences the relationship between sequencing depth and zero abundance (Figure 5a) [Hicks et al., 2015]. The large differences in sequencing depths between batches attenuate the overall association with zero abundance when cells are pooled across batches (Figure 5a). We therefore added a covariate to account for the batch effect in both the negative binomial mean (*μ*) and the zero inflation probability (*π*) of the ZINB-WaVE model used to produce the weights for DE analysis. Adjusting for batch yields weights with a slightly higher mode near zero, suggesting a more informative discrimination between excess and NB zeros (Figure 5b). Although the batch effect is small in terms of the weights, this illustrates the generality and flexibility of our ZINB-WaVE weighting approach: through a suitable parameterization of both the NB mean and ZI probability one can adjust for effects that can bias the weights and hence the DE results.

**Figure 5:**
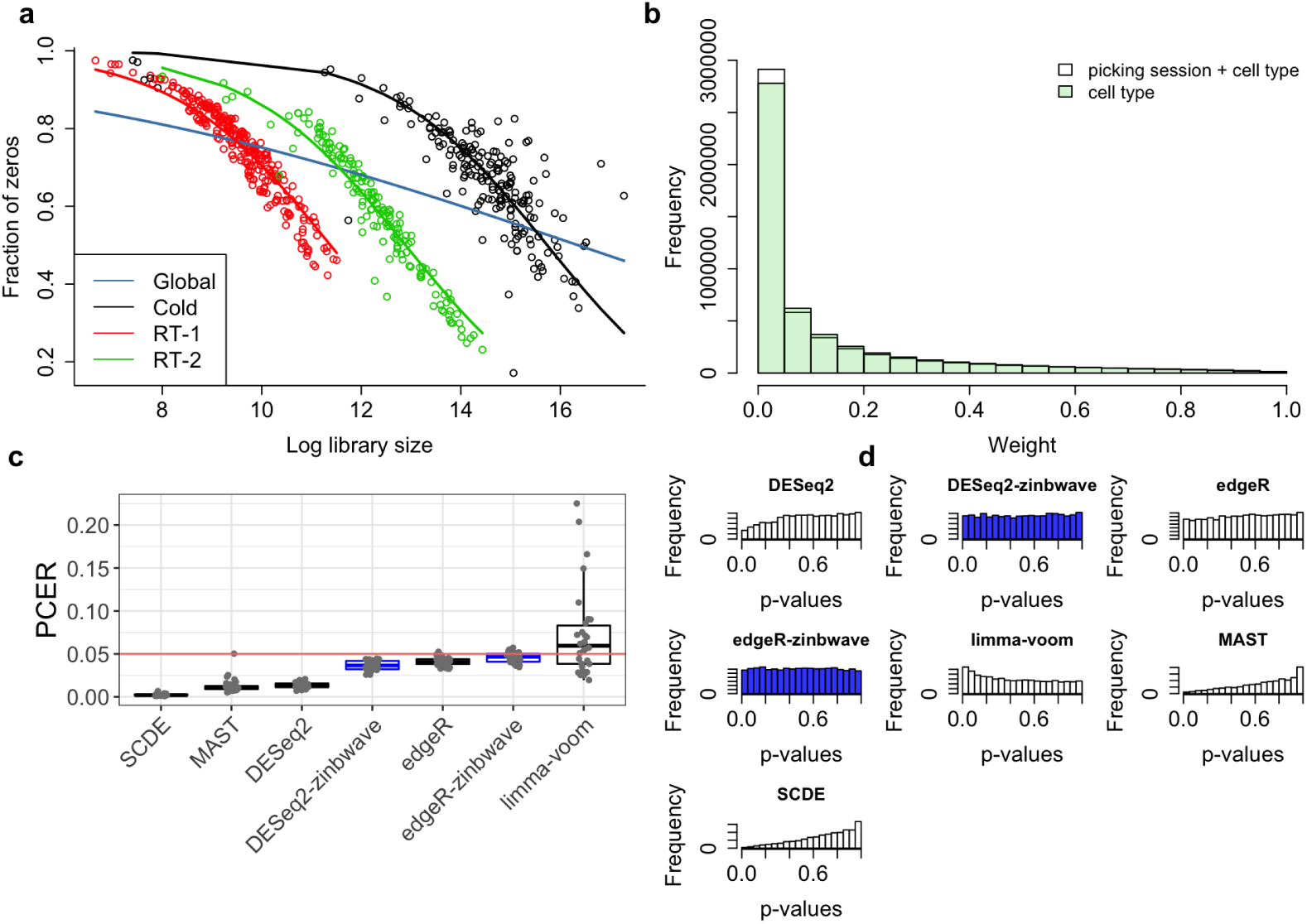
False positive control on mock null Usoskin datasets (*n* = 622 cells). (a) The scatterplot and GLM fits (R `glm` function with `family=binomial`), color-coded by batch (i.e., picking sessions Cold, RT-1, and RT-2), illustrate the association of zero abundance with sequencing depth. The three batches differ in their sequencing depths, causing an attenuated global relationship when pooling cells across batches (blue curve). Adjusting for the batch effect in the ZINB-WaVE model allows to properly account for the relationship between sequencing depth and zero abundance. (b) Histogram of ZINB-WaVE weights for zero counts for original Usoskin dataset, with (white) and without (green) including batch as a covariate in the ZINB-WaVE model. The higher mode near zero for batch adjustment indicates that more counts are being classified as dropouts, suggesting more informative discrimination between excess and NB zeros. (c) Boxplot of per-comparison error rate (PCER) for 30 mock null datasets for each of seven DE methods; ZINB-WaVE-weighted methods are highlighted in blue. (d) Histogram of unadjusted *p*-values for one of the datasets in (c). ZINB-WaVE was fit with intercept, cell type covariate (actual or mock), and batch covariate (unless specified otherwise) in *X*, *V* = 1_*J*_, *K* = 0 for *W*, common dispersion, and *ϵ* = 10^12^.

For the Usoskin dataset, we assessed false positive control by comparing the *actual* vs. the *nominal* PCER for mock null datasets where none of the genes are expected to be differentially expressed. Specifically, we generated 30 mock datasets where, for each dataset, two groups of 45 cells each were created by sampling 15 cells at random, without replacement from each of the three picking sessions. Sampling cells within batch allows to control for potential confounding by the batch variable. For each of the 30 mock datasets, we considered seven methods to identify genes that are DE between the two groups and declared a gene DE if its nominal unadjusted *p*-value was less than or equal to 0.05. For these mock datasets, any gene declared DE between the two groups is a false positive. Thus, for each method, the nominal PCER of 0.05 is compared to the actual PCER which is simply the proportion of genes declared DE (Figure 5c-d).

The seven methods considered are: unweighted and ZINB-WaVE-weighted `edgeR`, unweighted and ZINB-WaVE-weighted `DESeq2`, unweighted limma-voom (ZINB-WaVE-weighted limma-voom was found to perform poorly in the simulation study and hence is not considered here), `MAST`, and `SCDE` (see “Methods” for details). `edgeR` and `DESeq2` with ZINB-WaVE weights and unweighted `edgeR` controlled the PCER close to its nominal level (Figure 5c). The unweighted versions of `DESeq2`, `MAST`, and `SCDE` tended to be conservative, whereas limma-voom tended to be anti-conservative. In addition, the weighted versions of `edgeR` and `DESeq2` and unweighted `edgeR` yielded near uniform *p*-value distributions (as expected under this complete null scenario), while unweighted `DESeq2`, `MAST`, and `SCDE` tended to yield conservative *p*-values (mode near 1) and limma-voom anti-conservative *p*-values (mode near 0) (Figure 5d).

We also replicated the original analysis of Usoskin et al. [2014], by performing one-against-all tests of DE for each cell type (Additional File 1: Figure S13). limma-voom found a high number of DE genes, confirming our results from the mock evaluations where it was too liberal. The ZINB-WaVE methods tended to find a high number of DE genes, which is promising combined with the good PCER control seen in the mock comparisons. While introducing ZINB-WaVE weights in `DESeq2` lead to a higher number of significant genes on average, the effect is less clear with `edgeR` and seems to depend on the contrast.

Similar results were observed for a 10x Genomics PBMC dataset comprising 2, 700 single cells sequenced on the Illumina NextSeq 500 (Additional File 1: Figure S14), with the distinction that we found a conservative *p*-value distribution for ZINB-WaVE-weighted `DESeq2`. Since no information was provided about potential batch effects, we did not consider batch covariates for this dataset.

Additionally, we examined the PCER and *p*-value distributions on mock comparisons while varying the regularization parameter (*ϵ*) for the ZINB-WaVE estimation procedure. Not surprisingly, we observed that the PCER decreases with increasing *ϵ*, i.e., as the parameters of the ZINB-WaVE model are subjected to more “shrinking” (Additional File 1: Figures S15 and S16 for Usoskin and 10x Genomics PBMC datasets, respectively).

### Biologically meaningful clustering and differential expression results

To analyze the 2, 700 cells from the 10x Genomics PBMC dataset (see “Methods”), we followed the tutorial available at http://satijalab.org/seurat/pbmc3k_tutorial.html and used the R package Seurat [Butler and Satija, 2017]. The major steps of the pipeline were quality control, data filtering, identification of highvariance genes, dimensionality reduction using the first ten components from principal component analysis (PCA), and graph-based clustering. The final step of the pipeline was to identify genes that are differentially expressed between clusters, in order to derive cell type signatures. Two different parameterizations were used for the Seurat clustering. With one parameterization, a single cluster was identified for CD4+ T-cells, while with another, two CD4+ T-cell subclusters were identified, corresponding to CD4+ naive T-cells and CD4+ memory T-cells (gold and red clusters in Figure 6a, respectively). At the end of the tutorial, the authors concluded that the memory/naive split was weak and more cells would be needed to have a better separation between the two CD4+ T-cell subclusters.

**Figure 6:**
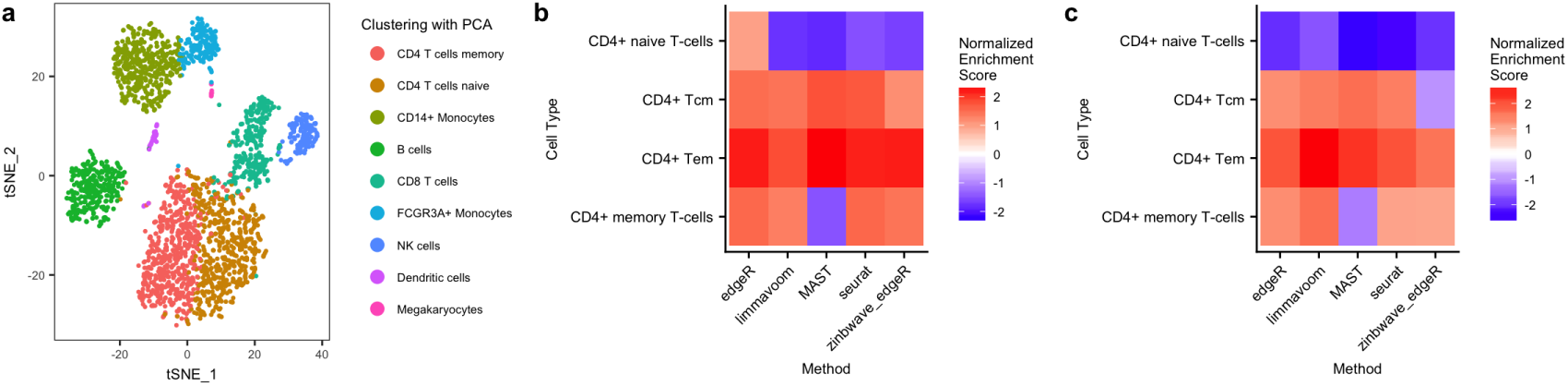
Biologically meaningful DE results for 10x Genomics PBMC dataset. (a) Scatterplot of the first two t-SNE dimensions obtained from the first 10 principal components. Cells are color-coded by clusters found using the Seurat graph-based clustering method on the first 10 principal components. Pseudo-color images on the right display normalized enrichment scores (NES) after gene set enrichment analysis (GSEA) for cell types related to CD4+ T-cells (see “Methods”), for clustering based on (b) the first 10 principal components and (c) *W* from ZINB-WaVE with *K* = 20. For dimensionality reduction, ZINB-WaVE was fit with *X* = 1_*n*_, *V* = 1_J_, *K* = 20 for *W* (based on AIC), common dispersion, and ϵ = 10^12^. To compute the weights for DE analysis, ZINB-WaVE was fit with intercept and cell type covariate in *X*, *V* = 1_*J*_, *K* = 0 for *W*, common dispersion, and *ε* = 10^12^. NES for more cell types are shown in Additional File 1: Figure S17.

In order to find DE genes between the two CD4+ T-cell subclusters, we used Seurat, unweighted `edgeR`, ZINB-WaVE-weighted `edgeR`, `MAST`, and limma-voom. We then sought to identify cell types using gene set enrichment analysis (GSEA), with the function fgsea from the Bioconductor R package fgsea [Sergushichev, 2016] and gene sets for 64 immune and stroma cell types from the R package ×Cell [Aran et al., 2017]. While unweighted `edgeR` found that one cluster was enriched in both CD4+ memory and naive T-cells compared to the other cluster, our weighted-edgeR method as well as Seurat, and limma-voom found that the cluster was enriched in CD4+ T-effector memory, CD4+ T-central memory, and CD4+ memory T-cells, and depleted in CD4+ naive T-cells. `MAST` found that the cluster was depleted in CD4+ memory T-cells and CD4+ naive T-cells, but enriched in CD4+ T-effector memory and CD4+ T-central memory T-cells (see Figure 6b and Additional File 1: Figure S17). This suggests that our ZINB-WaVE weights can successfully unlock `edgeR` for zero-inflated data, leading to biologically meaningful DE genes.

While ZINB-WaVE can be used to compute weights in a supervised setting with *a priori* known cell types, it can also be used to perform dimensionality reduction in an unsupervised setting. To demonstrate the ability of our method to find biologically relevant clusters and DE genes, we performed dimensionality reduction using ZINB-WaVE with K = 20 unknown covariates (matrix W, see “Methods”), where *K* = 20 was chosen using the Akaike information criterion (AIC) (Additional File 1: Figure S18). We then used W, instead of the first 10 components of PCA as in the Seurat tutorial, to cluster the cells using the Seurat graph-based clustering. We found similar clusters as the Seurat clusters, except for the NK-cell and B-cell clusters which were partitioned differently and the cluster with CD4+ T-cells (Additional File 1: Figure S19). Using this new clustering, GSEA showed a better separation between CD4+ naive T-cells and CD4+ memory T-cells for all the methods, suggesting a biological meaningful clustering using ZINB-WaVE dimensionality reduction instead of PCA. The CD4+ T-effector memory, CD4+ T-central memory, and CD4+ memory cell types were enriched using limma-voom, unweighted `edgeR`, `MAST`, and Seurat, but only the CD4+ T-central memory cell type was depleted using our weighted `edgeR` method (Figure 6c and Additional File 1: Figure S17). As we do not have prior knowledge about the cells in the different clusters, we are unable to say whether the cluster is more representative of the CD4+ T-effector memory cell type or if our method missed the enrichment in the CD4+ T-central memory cell type. However, it is interesting that using ZINB-WaVE to account for zero inflation in the clustering allowed `edgeR` to find results that seem more biologically meaningful than without accounting for zero inflation.

Finally, using a Benjamini and Hochberg [1995] adjusted *p*-value cut-off of 0.05, limma-voom declared 433 and 194 DE genes and weighted-edgeR 371 and 151, for clustering based on, respectively, the first 10 PCs and W from ZINB-WaVE. We additionally showed on mock comparisons for the same 10x Genomics PBMC dataset that limma-voom had a greater actual PCER than weighted `edgeR` (Additional File 1: Figure S14), suggesting that some of the DE genes found by limma-voom are likely to be false positives. This belief is reinforced by the skewed distribution of limma-voom *p*-values (Additional File 1: Figure S20).

### Alternative approaches to weight estimation

ZINB-WaVE is one particular approach to fit a ZINB model to single-cell RNA-seq data. However, our proposed data analysis strategy to unlock conventional RNA-seq tools with ZINB observation-level weights is not restricted to ZINB-WaVE-based workflows. In particular, we illustrate the use of weights estimated by the *zingeR* method, an expectation-maximization (EM) algorithm which we developed earlier and that builds upon `edgeR` for estimating the NB parameters of the ZINB model [Van den Berge et al., 2017]. The ZINB-WaVE and *zingeR* approaches differ in the following respects. The *zingeR* weights are based on a constant cell-specific excess zero probability *π*_*i*_ for each cell *i*, while the ZINB-WaVE excess zero probability *π*_*ij*_ is both cell and gene-specific, a strategy that was also advocated in recent methods [Pierson and Yau, 2015, Finak et al., 2015]. Secondly, the ZINB-WaVE negative binomial mean *μ* and zero inflation probability *π* are modeled in terms of both wanted and unwanted cell and gene-level covariates, allowing normalization for a variety of nuisance technical effects. Thirdly, different parameter estimation strategies are adopted: parameters from the *zingeR* model are estimated with an EM algorithm, whereas those from the ZINB-WaVE model are estimated using a penalized maximum likelihood approach. Finally, methods based on *zingeR* weights have the property of converging to their unweighted counterparts in the absence of zero inflation.

In terms of performance, based on the simulation study on full-length protocols, *zingeR* workflows dominate both bulk RNA-seq and dedicated scRNA-seq methods, but were found to be inferior in terms of sensitivity to ZINB-WaVE workflows (Additional File 1: Figure S21). However, for the Usoskin dataset, *zingeR* seems to find a higher number of DE genes than ZINB-WaVE and than its bulk RNA-seq counterparts (Additional File 1: Figure S22), while also controlling the PCER in mock evaluations (Additional File 1: Figure S23). Due to the computational burden of the *zingeR* method we were unable to apply it to large-scale datasets, such as those from the 10x Genomics platform, thus limiting our comparison.

### Computational time

The better performance of our ZINB-WaVE-weighted DE method comes at a computational cost, since we first fit ZINB-WaVE to the entire cells-by-genes matrix of read counts to compute the weights and then use a weighted version of `DESeq2` or `edgeR` for inferring DE. To give the reader an idea of how different methods scale in terms of computation time, we benchmarked three different datasets: the Islam dataset (92 cells), one of the mock null Usoskin datasets used in Figure 5 (90 cells), and the CD4+ T-cell cluster of the 10x Genomics PBMC dataset (1,151 cells). For each dataset, 10,000 genes were sampled at random and the two cell types were used as covariates. For the Usoskin dataset, batch was added as a covariate for all methods. For all datasets, the fastest method was limma-voom followed by `edgeR` (Additional File 1: Figure S24). As `DESeq2` was slower than `edgeR`, not surprisingly weighted-DESeq2 was also slower than weighted-edgeR, especially for the 10x Genomics PBMC dataset.

## Discussion

This manuscript focused on adapting standard bulk RNA-seq differential expression tools to handle the severe zero inflation present in single-cell RNA-seq data. We proposed a simple and general approach that integrates seamlessly with a range of popular DE software packages, such as `edgeR` and `DESeq2`. The main idea is to use weights for zero inflation in the negative binomial model underlying bulk RNA-seq methods. In particular, the weights are based on the ZINB-WaVE method of Risso et al. [2017]. The general and flexible ZINB-WaVE framework allows to extract low-dimensional signal from scRNA-seq read counts, accounting for zero inflation (e.g., dropouts), over-dispersion, and the discrete nature of the data. In particular, the ZINB-WaVE model allows for read count normalization through an appropriate parameterization of the negative binomial means and zero inflation probabilities in terms of both gene and cell-level covariates.

Our results complement the findings of Jaakkola et al. [2016] and Soneson and Robinson [2017a], that bespoke scRNA-seq tools do not systematically improve upon bulk RNA-seq tools. Although `MAST`, metagenomeSeq, and `SCDE` were explicitly developed to handle excess zeros, they suffer from poor performance in a high zero inflation setting, as demonstrated in the simulation study.

The value of our method was demonstrated for scRNA-seq protocols relying on both standard (Islam, Usoskin, and Trapnell datasets) and unique molecular identifier (UMI) (10x Genomics PBMC dataset) read counting. UMIs were recently proposed to reduce measurement variability across samples [Islam et al., 2013]. In UMI-based protocols, transcripts are labeled with a small random UMI barcode prior to amplification. After amplification and sequencing, one enumerates the unique UMIs found for every transcript, which correspond to individual sequenced UMI-labeled transcripts. There is some evidence in the literature that zero inflation is less of a problem for UMI-based than for full-length protocols and that UMI read counts could follow a negative binomial distribution [Grün et al., 2014, Ziegenhain et al., 2017]. Hence, our method also provides good results for UMI-based data with limited zero inflation, demonstrating its broad applicability.

In the simulation study, power to detect DE was generally lower for 10x Genomics UMI datasets (Figure 4) than for full-length protocol datasets (Figure 3). While the 10x Genomics platform has the advantage of an extremely high throughput, allowing many cells to be characterized, the resulting datasets often have the disadvantage of low library sizes, a logical consequence of UMI counting and of the trade-off between sequencing depth and number of cells to be sequenced in one sequencing run. As a result, the sequencing depth of these datasets is much lower than that of bulk RNA-seq datasets, making it harder to identify excess zeros and assess differential expression, even in large sample size settings. Although the 10x Genomics platform may be well suited for hypothesis generation, e.g., through cell type discovery or lineage trajectory studies, full-length protocols may be more appropriate for discovering marker genes between inferred cell types or trajectories, an approach that has also been adopted in previous studies [Pal et al., 2017].

We have used ZINB-WaVE in conjunction with either `edgeR` or `DESeq2`. However, the ZINB-WaVE posterior probabilities could be used as weights to unlock other standard RNA-seq workflows in zero inflation situations. Additional File 1: Figure S7 shows that ZINB-WaVE weights combined with heteroscedastic weights in limma-voom also increase power in a scRNA-seq context, although this may be at the expense of Type *i* error control.

The ZINB-WaVE method penalizes the L2 norm of the parameter estimates for regularization purposes. It requires a penalty parameter, *ϵ*, that is rescaled differently for gene-specific parameters, cell-specific parameters, and dispersion parameters [Risso et al., 2017]. All analyses in this manuscript were performed with *ϵ* = 10^12^, to provide consistently comparable results. However, the optimal value of *ϵ* is dataset-specific and further research is needed to provide a data-driven approach for selecting an optimal e. Indeed, based on our simulations, the value of the penalty parameter can have a profound influence on results (Additional File 1: Figure S25), but we found *ϵ* = 10^12^ to have generally good performance.

ZINB-WaVE allows the option to infer latent variables *W*, which may correspond to either unmeasured confounding covariates or unmeasured covariates of interest. The observational weights were computed with the number of unknown covariates *K* = 0, i.e., no latent variables were inferred. For clustering of the real datasets, we inferred an optimal choice of *K* using the AIC (Additional File 1: Figure S18). However, further investigation is needed to confirm that the AIC is appropriate for selecting *K*.

In principle, our proposed ZINB-WaVE model could also be used to identify DE genes both in terms of the negative binomial mean and the zero inflation probability, reflecting, respectively, a continuum in DE and a more binary (i.e., presence/absence) DE pattern. In this context, the parameters of interest are regression coefficients *β* corresponding to known sample-level covariates in the matrix *X* used in either *μ* or *π* (Equation (4)). Differentially expressed genes may be identified via likelihood ratio tests or Wald tests, with the standard errors of estimators of *β* obtained from the inverse of the Hessian matrix of the likelihood function. However, both types of tests would be computationally costly, as likelihood ratio tests would require refitting the entire model for each gene and Wald tests would require the Hessian matrix to be computed and inverted.

In this contribution, we have proposed to estimate the weights using ZINB-WaVE, but other approaches are possible. It is important to note that while methods such as ZINB-WaVE and *zingeR* can successfully identify excess zeros, they cannot however readily discriminate between their underlying causes, i.e., between technical (e.g., dropout) and biological (e.g., bursting) zeros. Although we cannot make this distinction with the weights, an increase in bursting rates between cell types, characterized by higher counts and more zeros [Fujita et al., 2016], can however be picked up by the count component of the ZINB model.

## Conclusions

In summary, we provide a realistic simulation framework for single-cell RNA-seq data and use the well-tested ZINB-WaVE method to successfully identify excess zeros and yield gene and cell-specific weights for differential expression analysis in scRNA-seq experiments. The tools we have developed allow an integrated workflow for normalization, dimensionality reduction, cell type discovery, and the identification of cell type marker genes. We confirmed that state-of-the-art scRNA-seq tools do not improve upon common bulk RNA-seq tools for differential expression analysis based on scRNA-seq data. Our workflow, however, outperforms current methods and has the merit of not deteriorating in performance in the absence of zero inflation. Inference of DE is focused on the count component of the ZINB model and our method produces posterior probabilities that can be used as observation-level weights by bulk RNA-seq tools. Hence, our approach unlocks widely-used bulk RNA-seq DE workflows for zero-inflated data and will assist researchers, data analysts, and developers in improving power to detect DE in the presence of excess zeros. The framework is general and applicable beyond scRNA-seq, to zero-inflated count data structures arising in applications such as metagenomics [Paulson et al., 2013, Xu et al., 2015].

## Methods

### ZINB-WaVE: Zero-inflated negative binomial-based wanted variation extraction

Zero-inflated distributions. A major difference between single-cell and bulk RNA-seq data is arguably the high abundance of zero counts in the former. Traditionally, excess zeros are dealt with by the use of hurdle or zero-inflated models, as recently proposed by Finak et al. [2015], Kharchenko et al. [2014], and Paulson et al. [2013]. A zero-inflated count distribution is a two-component mixture distribution between a point mass at zero and a count distribution, in our case, the negative binomial distribution which has been used successfully for bulk RNA-seq [Love et al., 2014, Robinson et al., 2010, Law et al., 2014, McCarthy et al., 2012b].

The probability mass function (PMF) *f*_*ZINB*_ for the zero-inflated negative binomial (ZINB) distribution is given by

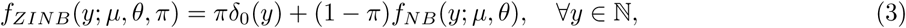

where *π ϵ* [0,1] denotes the mixture probability for zero inflation, *f*_*NB*_ (·; *μ*,*θ*) the negative binomial (NB) PMF with mean *μ* and dispersion *θ* = 1/*φ*, and *δ*_0_(·) the Dirac function (*δ*_0_(*y*) = when *y* = 0 and 0 otherwise and *δ*_0_ integrates to one over ℝ, i.e., has cumulative distribution function equal to *I*(*y* ≥ 0)). Here, *π* can be interpreted as the probability of an excess zero, i.e., inflated zero count, with respect to the NB distribution.

Under a ZINB model, the posterior probability that a given count y arises from the NB count component is given by Bayes’ rule

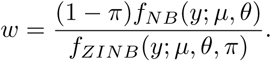

As described below, such posterior probabilities can be used as weights in standard bulk RNA-seq workflows, for a suitable parameterization of the ZI probability and NB mean.

#### ZINB-WaVE model

Given *n* observations (typically, *n* single cells) and *J* features (typically, *J* genes) that can be counted for each observation, let *Y*_*ij*_ denote the count of feature *j* (*j* = 1,…, *J*) for observation *i* (*i* = 1,…,*n*). To account for various technical and biological effects frequent in single-cell sequencing technologies, we model *Y*_*ij*_ as a random variable following a ZINB distribution with parameters *μ*_*ij*_, *θ*_*ij*_, and *π*_*ij*_, and consider the following regression models for these parameters:

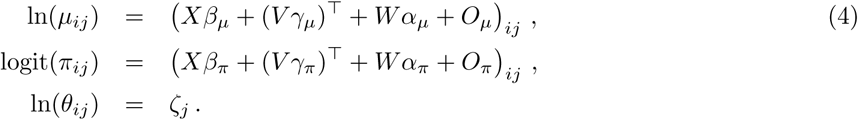

Both the NB mean expression level *μ* and the ZI probability *π* are modeled in terms of *observed sample-level and gene-level covariates* (*X* and *V*, respectively), as well as *unobserved sample-level covariates* (*W*) that need to be inferred from the data. *O*_*μ*_ and *O*_*π*_ are known matrices of offsets. The matrix *X* can include covariates that induce variation of interest, such as cell types, or covariates that induce unwanted variation, such as batch or quality control (QC) measures. It can also include a constant column of ones for an intercept that accounts for gene-specific global differences in mean expression level or dropout rate. The matrix *V* can include gene-level covariates, such as length or GC-content. It can also accommodate an intercept to account for cell-specific global effects, such as size factors representing differences in library sizes (i.e., total number of reads per sample). The unobserved matrix *W* contains unknown sample-level covariates, which could correspond to unwanted variation as in RUV [Gagnon-Bartsch and Speed, 2012, Risso et al., 2014] (e.g., *a priori* unknown batch effects) or could be of interest as in cluster analysis (e.g., *a priori* unknown cell types). The model extends the RUV framework to the ZINB distribution (thus far, RUV had only been implemented for linear [Gagnon-Bartsch and Speed, 2012] and log-linear regression [Risso et al., 2014]). It differs, however, in interpretation from RUV in the *W*_*α*_ term, which is not necessarily considered unwanted and generally refers to unknown low-dimensional variation. It is important to note that although *W* is the same, the matrices *X* and *V* could differ in the modeling of *μ* and *π*, if we assume that some known factors do not affect both.

As detailed in Risso et al. [2017], the model is fit using a penalized maximum likelihood estimation procedure.

### Using ZINB-WaVE weights in DE inference methods

We only consider statistical inference on the count component of the mixture distribution, that is, we are concerned with identifying genes whose expression levels are associated with covariates of interest as parameterized in the mean *μ* of the negative binomial component. Most popular bulk RNA-seq methods are based on the methodology of generalized linear models (GLM), which readily accommodates inference based on observation-level weights (R function `glm`), e.g., negative binomial model in Bioconductor R packages `edgeR` and `DESeq2`. Note that although the ZINB-WaVE weights are gene and cell-specific, the GLMs are fit gene by gene; hence, for given gene, the cell-specific weights are used as observation-specific weights in the GLMs.

#### edgeR

We extended the `edgeR` package [Robinson et al., 2010, McCarthy et al., 2012b] by fitting a negative binomial model genewise, with ZINB-WaVE posterior probabilities as observation-level weights in the weights slot of an object of class *DGEList*, and estimating the dispersion parameter by the usual approximate empirical Bayes shrinkage. Downweighting is accounted for by adjusting the degrees of freedom of the null distribution of the test statistics. Specifically, we reintroduced the moderated *F*-test from a previous version of `edgeR`, where the denominator residual degrees of freedom *df*_*j*_ for a particular gene *j* are adjusted by the extent of zero inflation identified for this gene, i.e., *df*_*j*_ = ∑*_i_ w*_*ij*_ – *p*, where *w*_*ij*_ is the ZINB-WaVE weight for gene *j* in cell *i* and *p* the number of parameters estimated in the NB generalized linear model. This weighted *F*-test is implemented in the function `glmWeightedF` from the Bioconductor R package zinbwave.

#### DESeq2

We extended the `DESeq2` package [Love et al., 2014] to accommodate zero inflation by providing the option to use observation-level weights in the parameter estimation steps. In this case, the ZINB-WaVE weights are supplied in the weight slot of an object of class *DESeqDataSet*.

`DESeq2`’s default normalization procedure is based on geometric means of counts, which are zero for genes with at least one zero count. This greatly limits the number of genes that can be used for normalization in scRNA-seq applications [Vallejos et al., 2017]. We therefore use the normalization method suggested in the phyloseq package [McMurdie and Holmes, 2013], which calculates the geometric mean for a gene by only using its positive counts, so that genes with zero counts could still be used for normalization purposes. The phyloseq normalization procedure can now be applied by setting the argument `type` equal to `poscounts` in the `DESeq2` function `estimateSizeFactors`. For single-cell UMI data, for which the expected counts may be very low, the likelihood ratio test implemented in `nbinomLRT` should be used. For other protocols (i.e., non-UMI), the Wald test in `nbinomWaldTest` can be used, with null distribution a *t*-distribution with degrees of freedom corrected for downweighting. In both cases, we recommend the minimum expected count to be set to a small value (`minmu=1e-6`). The Wald test in `DESeq2` allows for testing contrasts of the coefficients.

#### limma-voom

For the limma-voom approach [Law et al., 2014], implemented in the voom function from the limma package, heteroscedastic weights are estimated based on the mean-variance relationship of the log-transformed counts. We adapt limma-voom to zero-inflated situations by multiplying the heteroscedastic weights by the ZINB-WaVE weights and using the resulting weights in weighted linear regression. To account for the downweighting of zeros, the residual degrees of freedom of the linear model are adjusted as with `edgeR` before the empirical Bayes variance shrinkage and are therefore also propagated to the moderated statistical tests. Both the standard and ZINB-WaVE-weighted versions of limma-voom were considered in the simulation study; the latter was not considered for the real datasets, due to its poor performance in the simulation study.

#### Multiple testing

For the simulation study, in order to reduce the number of tests performed [Bourgon et al., 2010], we apply the independent filtering procedure implemented in the genefilter package and used in `DESeq2` [Love et al., 2014]. As in `DESeq2`, we exclude from the multiple testing correction any gene whose average expression strength (i.e., average of fitted values) is below a threshold chosen to maximize the number of differentially expressed genes. Note that the filtering procedure can affect each method differently, due to differences in fitted values and *p*-value distributions.

Unless specified otherwise, the *p*-values for all methods are then adjusted using the Benjamini and Hochberg [1995] procedure for controlling the false discovery rate (FDR).

#### Performance assessment

We assess performance based on scatterplots of the true positive rate (TPR) vs. the false discovery proportion (FDP), as well as receiver operating characteristic (ROC) curves of the true positive rate (TPR) vs. the false positive rate (FPR), according to the following definitions

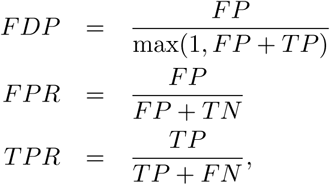

where *FN, FP, TN*, and *TP* denote, respectively, the numbers of false negative, false positives, true negatives, and true positives. FDP-TPR curves and ROC curves are implemented in the Bioconductor R package iCOBRA [Soneson and Robinson, 2016].

#### DE method comparison

We compared our weighted DE approach to state-of-the-art bulk RNA-seq methods implemented in the packages `edgeR` (v3.20.1) [Robinson et al., 2010, McCarthy et al., 2012b], `DESeq2` (v1.19.8) [Love et al., 2014], and limma (v3.34.0) [Law et al., 2014]. We also considered dedicated scRNA-seq tools from the packages `scde` (v2.6.0) [Kharchenko et al., 2014], `MAST` (v1.4.0) [Finak et al., 2015], and NODES (v0.0.0.9010) [Sengupta et al., 2016], as well as metagenomeSeq (v1.18.0) [Paulson et al., 2013] developed to account for zero inflation in metagenomics applications. A ZINB model is also implemented in ShrinkBayes [van de Wiel et al., 2014], but the method does not scale to the typical sample sizes encountered in scRNA-seq and has many tuning parameters, which lead us to not include it in our comparison. In `DESeq2`, we disable the outlier imputation step and allow for shrinkage of fold-changes by default. In addition, for large 3′-end sequencing datasets like the Usoskin and 10x Genomics PBMC datasets, we set the minimum expected count estimated by `DESeq2` to 10^‒6^, allowing the method to cope with large sample sizes and low counts. We use the recommended gene filtering procedures for NODES and `MAST`, except for computing time benchmarking, where no genes are filtered out to allow a fair comparison. For all other methods, arguments were set to their default values.

### scRNA-seq data simulation

We extended the framework of Zhou et al. [2014] towards scRNA-seq applications and provide user-friendly R code to simulate scRNA-seq read counts in the GitHub repository linked to this manuscript (https://github.com/statOmics/zinbwaveZinger). The user can input a real scRNA-seq dataset to infer gene-level parameters for the read count distributions. Library sizes for the simulated samples are by default resampled from the real dataset, but can also be user-specified. The simulation paradigm randomly resamples parameters estimated from the original dataset, where all parameters of a given gene are resampled jointly in order to retain gene-specific characteristics present in the original dataset.

In scRNA-seq, dropouts and bursting lead to bias in parameter estimation. Our simulation framework alleviates this problem by using zero-truncated negative binomial (ZTNB) method-of-moments estimators [Moore, 1986, McCullagh and Nelder, 1989] on the positive counts to estimate the expression fraction *λ*_*j*_ = *E*[*Y*_*ij*_/*N*_*i*_], with *N*_*i*_ = ∑_*j*_ *Y*_*ij*_ the sequencing depth of cell *i*, and the negative binomial dispersion *θ*_*j*_ = 1/*φ*_*j*_. Specifically, initial NB-based estimators are iteratively updated according to the ZTNB-based estimators provided by

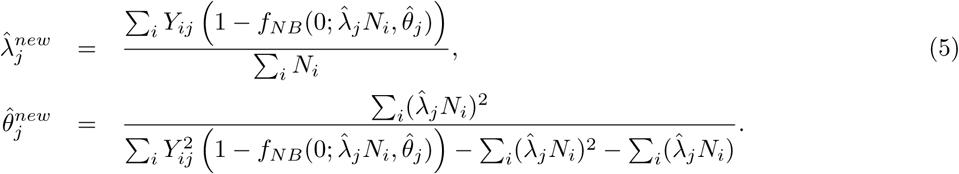

Note that, when *Y*_*ij*_ is zero, it does not contribute to the estimators of *λ*_*j*_ and *θ*_*j*_. These estimates are then used to simulate counts according to a negative binomial distribution.

We additionally simulate excess zeros by modeling the empirical zero abundance *p*_*ij*_ = *I*(*Y*_*ij*_ = 0) as a function of an interaction between the gene-specific expression intensity, measured as average log count per million (CPM)

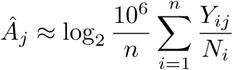

(as calculated using the `aveLogCPM` function from `edgeR`), and the cell-specific sequencing depth *N*_*i*_, using a semi-parametric additive logistic regression model,

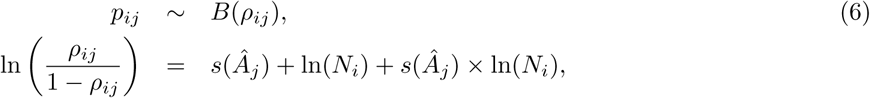

where *B*(*ρ*_*ij*_) denotes the Bernoulli distribution with parameter *ρ*_*ij*_ and *s*(·) a non-parametric thin-plate spline [Wood, 2003]. We then compare, for every gene, the estimated probability of zero counts based on the model in Equation (6) to the corresponding NB-based probability *f*_*NB*_ (0; *μ̂*_*ij*_, *θ̂*_*j*_) with *μ̂*_*ij*_ = *λ̂*_*j*_*N*_*i*_, and randomly add excess zeros whenever the former probability is higher than the latter. The model in Equation (6) is motivated by dataset-specific associations observed in real scRNA-seq datasets (Additional File 1: Figures S4–S5).

This framework acknowledges both gene-specific characteristics as well as broad dataset-specific associations across all genes and provides realistic scRNA-seq data for method evaluation. We assessed performance of various DE methods using data simulated based on the Islam et al. [2011] dataset, a subset of the Trapnell et al. [2013] dataset, and a 10x Genomics PBMC dataset. See the “Real datasets” section for information on these datasets.

### Gene set enrichment analysis

To identify cell types corresponding to the two CD4+ T-cell subclusters of the 10x Genomics PBMC dataset, we used gene set enrichment analysis (GSEA) with the function fgsea from the Bioconductor R package fgsea (v1.4.0) [Sergushichev, 2016] and gene sets for 64 immune and stroma cell types from the R package ×Cell (v1.1.0) [Aran et al., 2017]. For each DE method, the input to fgsea is a list of genes ranked by a test statistic comparing expression in the two CD4+ T-cell subclusters.

To facilitate comparison between DE methods, the test statistic used here is a transformation of the unadjusted *p*-values (*p*) with the sign of the log-fold-change (*lfc*): Φ^‒1^(1 – *p*/2) * sign(*lfc*), where Φ(·) denotes the standard Gaussian cumulative distribution function. As suggested by fgsea, all genes were used for the analysis. To assess the enrichment/depletion of one cluster compared to the other cluster, we used the normalized enrichment score (NES). The enrichment score (ES) is the same as in the Broad GSEA implementation [Subramanian et al., 2005] and reflects the degree to which a gene set is overrepresented at the top or bottom of a ranked list of genes. Briefly, the ES is calculated by walking down the ranked list of genes, increasing a running-sum statistic when a gene is in the gene set and decreasing it when it is not. A positive ES indicates enrichment at the top of the ranked list; a negative ES indicates enrichment at the bottom of the ranked list. The enrichment score is then normalized by the mean enrichment of random samples of genes, where genes are permuted from the original ranked list (10,000 permutations were used).

### Real datasets

#### Usoskin dataset

This dataset concerns mouse neuronal cells from the dorsal root ganglion, sequenced on either an Illumina Genome Analyzer IIx or HiSeq 2000 [Usoskin et al., 2014]. The cells were robotically picked in three separate sessions and the 5′-end of the transcripts sequenced. The expression measures were downloaded from “Supplementary Data” accompanying the original manuscript (http://linnarssonlab.org/drg/. After quality control and sample filtering (removal of non-single cells and non-neuronal cells), the authors considered 622 cells which were classified into eleven neuronal cell type categories. Only the genes with more than 20 non-zero counts were retained, for a total of 12, 132 genes.

The authors acknowledge the existence of a batch effect related to the picking session for the cells. For differential expression analysis, the picking session was therefore included as a batch covariate in all models.

To mimic a null dataset with no differential expression, we created two groups of 45 cells each, where, for each group, 15 cells were sampled at random, without replacement (over all cell types) from each picking session. For each of 30 such mock null datasets, we considered seven methods to identify genes that are DE between the two groups and declared a gene DE if its nominal unadjusted *p*-value was less than or equal to 0.05. For these mock datasets, any gene declared DE between the two groups is a false positive. Thus, the *nominal* per-comparison error rate (PCER) of 0.05 for each method is compared to its *actual* PCER which is simply the proportion of genes declared DE.

#### 10x Genomics PBMC dataset

We analyzed a dataset of peripheral blood mononuclear cells (PBMC) freely available from 10x Genomics (https://support.10xgenomics.com/single-cell-gene-expression/datasets/1.1.0/pbmc3k). We downloaded the data from https://s3-us-west-2.amazonaws.com/10x.files/samples/cell/pbmc3k/pbmc3k_filtered_gene_bc_matrices.tar.gz, which correspond to 2,700 single cells sequenced on the Illumina NextSeq 500 using unique molecular identifiers (UMI). We clustered cells following the tutorial available at http://satijalab.org/seurat/pbmc3k_tutorial.html and using the R package Seurat (v2.1.0) [Butler and Satija, 2017]. The major steps of the pipeline are quality control, data filtering, identification of high-variance genes, dimensionality reduction using the first ten components from principal component analysis (PCA), and graph-based clustering. To identify cluster markers, we used our ZINB-WaVE-weighted DE method instead of the method implemented in `Seurat`.

We created 30 mock null datasets and identified DE genes on these as for the Usoskin dataset, i.e., we created two groups of 45 cells each, by sampling at random, without replacement from the 2, 700 cells of the real dataset (no batch information available).

#### Islam dataset

The count table for the Islam et al. [2011] dataset was downloaded from the Gene Expression Omnibus (GEO) with accession number GSE29087. The Islam dataset represents 44 embryonic fibroblasts and 48 embryonic stem cells in the mouse, sequenced on an Illumina Genome Analyzer II. Negative control wells were removed and only the 11, 796 genes with at least 5 positive counts were retained for analysis. For the simulation, we generated datasets with 2 groups of 40 cells each.

#### Trapnell dataset

The dataset from Trapnell et al. [2013] was downloaded from the preprocessed single-cell data repository conquer (http://imlspenticton.uzh.ch:3838/conquer). Cells were sequenced on either an Illumina HiSeq 2000 or HiSeq 2500. We only used the subset of cells corresponding to the 48h and 72h timepoints of differentiating human myoblasts in order to generate two-group comparisons. Wells that do not contain one cell or that contain debris were removed. We used a more stringent gene filtering criterion than for the Islam dataset and retained the 24, 576 genes with at least 10 positive counts. The simulated datasets contain two conditions with 75 cells in each condition, thereby replicating the sample sizes of the Trapnell dataset.

### Software implementation

An R software package for our novel scRNA-seq simulation framework is available on the GitHub repository for this manuscript (https://github.com/statOmics/zinbwaveZinger) and is soon to be submitted to the Bioconductor project (http://www.bioconductor.org). The ZINB-WaVE weight computation is implemented in the computeObservationalWeights function of the Bioconductor R package zinbwave. ZINB-WaVE-weighted `edgeR` can be implemented using the glmWeightedF function from the zinb-wave package, while ZINB-WaVE-weighted `DESeq2` can be implemented using the native nbinomWaldTest function from the `DESeq2` package. More details on a ZINB-WaVE-weighted analysis can be found in the zinbwave vignette (http://bioconductor.org/packages/zinbwave/). Additionally, all analyses and figures reported in the manuscript can be reproduced using code provided in the GitHub repository (https://github.com/statOmics/zinbwaveZinger.

## Declarations

### Funding

This research was supported in part by IAP research network “StUDyS” grant no. P7/06 of the Belgian government (Belgian Science Policy) and the Multidisciplinary Research Partnership “Bioinformatics: from nucleotides to networks” of Ghent University. KVDB is supported by a Strategic Basic Research PhD grant from the Research Foundation - Flanders (FWO) no. 1S 418 16N. CS is supported by the Forschungskredit of the University of Zurich, grant no. FK-16-107. ML is supported by NIH grant no. CA142538-08. DR and SD were supported by the National Institutes of Health BRAIN Initiative (grant U01 MH105979, PI: John Ngai). JPV was supported by the French National Research Agency (grant ABS4NGS ANR-11-BINF-0001), the European Research Council (grant ERC-SMAC-280032), the Miller Institute for Basic Research in Science, and the Fulbright Foundation.

### Authors’ contributions

KVDB, FP, DR, SD, and LC conceived the methodology and designed the study, with input from all other authors. KVDB, FP, DR, and LC implemented the method and KVDB and FP performed the analyses. ML extended the `DESeq2` package. KVDB, FP, SD, and LC wrote the manuscript. All authors read and approved the final manuscript.

## Acknowledgements

We thank Tine Descamps and the students of the Statistical Genomics course, 2015-2016, Ghent University, who assisted us in assessing our initial implementation to unlock RNA-seq tools for zero inflation during `MAST`er thesis and project work, respectively.

## Additional Files

Additional file 1 - Supplementary figures

Additional file 2 - `countsimQC` evaluation of simulated Islam dataset.

Additional file 3 - `countsimQC` evaluation of simulated Trapnell dataset.

Additional file 4 - `countsimQC` evaluation of simulated 10X dataset.

## References

1. Michael I Love, Wolfgang Huber, and Simon Anders. Moderated estimation of fold change and dispersion for RNA-seq data with DESeq2. Genome Biology, 15(12):550, dec 2014. ISSN 1465-6906. doi: 10.1186/s13059-014-0550-8. URL http://genomebiology.com/2014/15/12/550.

2. Mark D Robinson, Davis J McCarthy, and Gordon K Smyth. edgeR: a Bioconductor package for differential expression analysis of digital gene expression data. Bioinformatics (Oxford, England), 26(1):139–40, jan 2010. ISSN 1367-4811. doi: 10.1093/bioinformatics/btp616. URL http://www.ncbi.nlm.nih.gov/pubmed/19910308http://www.pubmedcentral.nih.gov/articlerender.fcgi?artid=PMC2796818.

3. Charity W Law, Yunshun Chen, Wei Shi, and Gordon K Smyth. voom: Precision weights unlock linear model analysis tools for RNA-seq read counts. Genome biology, 15(2):R29, jan 2014. ISSN 1465-6914. doi: 10.1186/gb-2014-15-2-r29. URL http://www.pubmedcentral.nih.gov/articlerender.fcgi?artid=4053721{&}tool=pmcentrez{&}rendertype=abstract.

4. Zhong Wang, Mark Gerstein, and Michael Snyder. RNA-Seq: a revolutionary tool for transcriptomics. Nature Reviews Genetics, 10(1):57–63, jan 2009. ISSN 1471-0056. doi: 10.1038/nrg2484. URL http://www.nature.com/doifinder/10.1038/nrg2484.

5. Sara Goodwin, John D. McPherson, and W. Richard McCombie. Coming of age: ten years of next-generation sequencing technologies. Nature Reviews Genetics, 17(6):333–351, may 2016. ISSN 1471-0056. doi: 10. 1038/nrg.2016.49. URL http://www.nature.com/doifinder/10.1038/nrg.2016.49.

6. Tapio Lonnberg, Valentine Svensson, Kylie R James, Daniel Fernandez-Ruiz, Ismail Sebina, Ruddy Montandon, Megan S F Soon, Lily G Fogg, Arya Sheela Nair, Urijah Liligeto, Michael J T Stubbington, Lam-Ha Ly, Frederik Otzen Bagger, Max Zwiessele, Neil D Lawrence, Fernando Souza-Fonseca-Guimaraes, Patrick T Bunn, Christian R Engwerda, William R Heath, Oliver Billker, Oliver Stegle, Ashraful Haque, and Sarah A Teichmann. Single-cell RNA-seq and computational analysis using temporal mixture modelling resolves Th1/Tfh fate bifurcation in malaria. Science immunology, 2(9), mar 2017. doi: 10.1126/sciimmunol.aal2192. URL http://www.ncbi.nlm.nih.gov/pubmed/28345074http://www.pubmedcentral.nih.gov/articlerender.fcgi?artid=PMC5365145.

7. Florian Buettner, Kedar N Natarajan, F Paolo Casale, Valentina Proserpio, Antonio Scialdone, Fabian J Theis, Sarah A Teichmann, John C Marioni, and Oliver Stegle. Computational analysis of cell-to-cell heterogeneity in single-cell RNA-sequencing data reveals hidden subpopulations of cells. Nature Biotechnology, 33(2):155–160, jan 2015. ISSN 1087-0156. doi: 10.1038/nbt.3102. URL http://www.ncbi.nlm.nih.gov/pubmed/25599176http://www.nature.com/doifinder/10.1038/nbt.3102.

8. Anoop P. Patel, Itay Tirosh, John J. Trombetta, Alex K. Shalek, Shawn M. Gillespie, Hiroaki Wakimoto, Daniel P. Cahill, Brian V. Nahed, William T. Curry, Robert L. Martuza, David N. Louis, Orit Rozenblatt-Rosen, Mario L. Suva, Aviv Regev, and Bradley E. Bernstein. Single-cell RNA-seq highlights intratumoral heterogeneity in primary glioblastoma. Science, 344(6190), 2014. URL http://science.sciencemag.org/content/344/6190/1396.

9. Aleksandra A Kolodziejczyk, Jong Kyoung Kim, Jason C H Tsang, Tomislav Ilicic, Johan Henriksson, Kedar N Natarajan, Alex C Tuck, Xuefei Gao, Marc Bühler, Pentao Liu, John C Marioni, and Sarah A Teichmann. Single Cell RNA-Sequencing of Pluripotent States Unlocks Modular Transcriptional Variation. Cell stem cell, 17(4):471–85, oct 2015a. ISSN 1875-9777. doi: 10.1016/j.stem.2015. 09.011. URL http://www.ncbi.nlm.nih.gov/pubmed/26431182http://www.pubmedcentral.nih.gov/articlerender.fcgi?artid=PMC4595712.

10. Li Li, Ji Dong, Liying Yan, Jun Yong, Xixi Liu, Yuqiong Hu, Xiaoying Fan, Xinglong Wu, Hongshan Guo, Xiaoye Wang, Xiaohui Zhu, Rong Li, Jie Yan, Yuan Wei, Yangyu Zhao, Wei Wang, Yixin Ren, Peng Yuan, Zhiqiang Yan, Boqiang Hu, Fan Guo, Lu Wen, Fuchou Tang, and Jie Qiao. Single-Cell RNA-Seq Analysis Maps Development of Human Germline Cells and Gonadal Niche Interactions. Cell Stem Cell, 20 (6):858–873.e4, jun 2017. ISSN 19345909. doi: 10.1016/j.stem.2017.03.007. URL http://www.ncbi.nlm.nih.gov/pubmed/28457750http://linkinghub.elsevier.com/retrieve/pii/S1934590917300784.

11. Dmitry Usoskin, Alessandro Furlan, Saiful Islam, Hind Abdo, Peter Lönnerberg, Daohua Lou, Jens Hjerling-Leffler, Jesper Haeggström, Olga Kharchenko, Peter V Kharchenko, Sten Linnarsson, and Patrik Ernfors. Unbiased classification of sensory neuron types by large-scale single-cell RNA sequencing. Nature Neuroscience, 18(1):145–153, nov 2014. ISSN 1097-6256. doi: 10.1038/nn.3881. URL http://www.nature.com/doifinder/10.1038/nn.3881.

12. Aleksandra A. Kolodziejczyk, Jong Kyoung Kim, Valentine Svensson, John C. Marioni, and Sarah A. Teichmann. The Technology and Biology of Single-Cell RNA Sequencing. Molecular Cell, 58(4):610–620, may 2015b. ISSN 10972765. doi: 10.1016/j.molcel.2015.04.005. URL http://linkinghub.elsevier.com/retrieve/pii/S1097276515002610.

13. T. Nakamura, Y. Yabuta, I. Okamoto, S. Aramaki, S. Yokobayashi, K. Kurimoto, K. Sekiguchi, M. Nakagawa, T. Yamamoto, and M. Saitou. SC3-seq: a method for highly parallel and quantitative measurement of single-cell gene expression. Nucleic Acids Research, 43(9):e60–e60, may 2015. ISSN 0305-1048. doi: 10.1093/nar/gkv134. URL https://academic.oup.com/nar/article-lookup/doi/10.1093/nar/gkv134.

14. Angela R Wu, Norma F Neff, Tomer Kalisky, Piero Dalerba, Barbara Treutlein, Michael E Rothenberg, Francis M Mburu, Gary L Mantalas, Sopheak Sim, Michael F Clarke, and Stephen R Quake. Quantitative assessment of single-cell RNA-sequencing methods. Nature Methods, 11(1):41–46, oct 2013. ISSN 1548-7091. doi: 10.1038/nmeth.2694. URL http://www.ncbi.nlm.nih.gov/pubmed/24141493http://www.pubmedcentral.nih.gov/articlerender.fcgi?artid=PMC4022966http://www.nature.com/doifinder/10.1038/nmeth.2694.

15. Saiful Islam, Amit Zeisel, Simon Joost, Gioele La Manno, Pawel Zajac, Maria Kasper, Peter Lönnerberg, and Sten Linnarsson. Quantitative single-cell RNA-seq with unique molecular identifiers. Nature Methods, 11 (2):163–166, dec 2013. ISSN 1548-7091. doi: 10.1038/nmeth.2772. URL http://www.ncbi.nlm.nih.gov/pubmed/24363023http://www.nature.com/doifinder/10.1038/nmeth.2772.

16. Saiful Islam, Una Kjällquist, Annalena Moliner, Pawel Zajac, Jian-Bing Fan, Peter Lönnerberg, and Sten Linnarsson. Characterization of the single-cell transcriptional landscape by highly multiplex RNA-seq. Genome research, 21(7):1160–7, jul 2011. ISSN 1549-5469. doi: 10.1101/gr.110882.110. URL http://www.pubmedcentral.nih.gov/articlerender.fcgi?artid=3129258{&}tool=pmcentrez{&}rendertype=abstract.

17. Simone Picelli, Omid R Faridani, ?sa K Bj?rklund, G?sta Winberg, Sven Sagasser, and Rickard Sandberg. Full-length RNA-seq from single cells using Smart-seq2. Nature Protocols, 9(1):171–181, jan 2014. ISSN 1754-2189. doi: 10.1038/nprot.2014.006. URL http://www.ncbi.nlm.nih.gov/pubmed/24385147http://www.nature.com/doifinder/10.1038/nprot.2014.006.

18. Tamar Hashimshony, Naftalie Senderovich, Gal Avital, Agnes Klochendler, Yaron de Leeuw, Leon Anavy, Dave Gennert, Shuqiang Li, Kenneth J Livak, Orit Rozenblatt-Rosen, Yuval Dor, Aviv Regev, and Itai Yanai. CEL-Seq2: sensitive highly-multiplexed single-cell RNA-Seq. Genome biology, 17:77, apr 2016. ISSN 1474-760X. doi: 10.1186/s13059-016-0938-8. URL http://www.ncbi.nlm.nih.gov/pubmed/27121950http://www.pubmedcentral.nih.gov/articlerender.fcgi?artid=PMC4848782.

19. Greg Finak, Andrew McDavid, Masanao Yajima, Jingyuan Deng, Vivian Gersuk, Alex K. Shalek, Chloe K. Slichter, Hannah W. Miller, M. Juliana McElrath, Martin Prlic, Peter S. Linsley, and Raphael Gottardo. MAST: a flexible statistical framework for assessing transcriptional changes and characterizing heterogeneity in single-cell RNA sequencing data. Genome Biology, 16(1):278, dec 2015. ISSN 1474-760X. doi: 10.1186/s13059-015-0844-5. URL http://genomebiology.com/2015/16/1/278.

20. Arjun Raj and Alexander van Oudenaarden. Nature, nurture, or chance: stochastic gene expression and its consequences. Cell, 135(2):216–26, oct 2008. ISSN 1097-4172. doi: 10.1016/j.cell.2008.09.050. URL http://www.ncbi.nlm.nih.gov/pubmed/18957198http://www.pubmedcentral.nih.gov/articlerender.fcgi?artid=PMC3118044.

21. Arjun Raj, Charles S Peskin, Daniel Tranchina, Diana Y Vargas, and Sanjay Tyagi. Stochastic mRNA Synthesis in Mammalian Cells. PLoS Biology, 4(10):e309, sep 2006. ISSN 1545-7885. doi: 10.1371/journal.pbio.0040309. URL http://www.ncbi.nlm.nih.gOv/pubmed/17048983http://www.pubmedcentral.nih.gov/articlerender.fcgi?artid=PMC1563489http://dx.plos.org/10.1371/journal.pbio.0040309.

22. Emma Pierson and Christopher Yau. ZIFA: Dimensionality reduction for zero-inflated single-cell gene expression analysis. Genome Biology 2015 16:1, 16(1):241, nov 2015. ISSN 1474-760X. doi: 10.1186/s13059-015-0805-z. URL https://genomebiology.biomedcentral.com/articles/10.1186/s13059-015-0805-z.

23. Davide Risso, Fanny Perraudeau, Svetlana Gribkova, Sandrine Dudoit, and Jean-Philippe Vert. ZINB-WaVE: A general and flexible method for signal extraction from single-cell RNA-seq data. bioRxiv, 2017. doi: 10.1101/125112. URL http://www.biorxiv.org/content/early/2017/04/06/125112.

24. Manu Setty, Michelle D Tadmor, Shlomit Reich-Zeliger, Omer Angel, Tomer Meir Salame, Pooja Kathail, Kristy Choi, Sean Bendall, Nir Friedman, and Dana Pe’er. Wishbone identifies bifurcating developmental trajectories from single-cell data. Nature biotechnology, 34(6):637–45, jun 2016. ISSN 1546-1696. doi: 10.1038/nbt.3569. URL http://www.ncbi.nlm.nih.goV/pubmed/27136076http://www.pubmedcentral.nih.gov/articlerender.fcgi?artid=PMC4900897.

25. Xiaojie Qiu, Qi Mao, Ying Tang, Li Wang, Raghav Chawla, Hannah A Pliner, and Cole Trapnell. Reversed graph embedding resolves complex single-cell trajectories. Nature Methods, aug 2017. doi: 10.1038/nmeth. 4402. URL https://www.nature.com/nmeth/journal/vaop/ncurrent/full/nmeth.4402.html.

26. Kelly Street, Davide Risso, Russell B Fletcher, Diya Das, John Ngai, Nir Yosef, Elizabeth Purdom, and Sandrine Dudoit. Slingshot: Cell lineage and pseudotime inference for single-cell transcriptomics. bioRxiv, page 128843, apr 2017. doi: 10.1101/128843. URL http://www.biorxiv.org/content/early/2017/04/19/128843.

27. Aaron T.L. Lun, Davis J. McCarthy, and John C. Marioni. A step-by-step workflow for low-level analysis of single-cell RNA-seq data with Bioconductor. F1000Research, 5:2122, oct 2016. ISSN 2046-1402. doi: 10.12688/f1000research.9501.2. URL https://f1000research.com/articles/5-2122/v2.

28. Peter V Kharchenko, Lev Silberstein, and David T Scadden. Bayesian approach to single-cell differential expression analysis. Nature methods, 11(7):740–2, jul 2014. ISSN 1548-7105. doi: 10.1038/ nmeth.2967. URL http://www.ncbi.nlm.nih.gov/pubmed/24836921http://www.pubmedcentral.nih.gov/articlerender.fcgi?artid=PMC4112276.

29. Maria K. Jaakkola, Fatemeh Seyednasrollah, Arfa Mehmood, and Laura L. Elo. Comparison of methods to detect differentially expressed genes between single-cell populations. Briefings in Bioinformatics, page bbw057, jul 2016. ISSN 1467-5463. doi: 10.1093/bib/bbw057. URL http://www.ncbi.nlm.nih.gov/pubmed/27373736http://bib.oxfordjournals.org/lookup/doi/10.1093/bib/bbw057.

30. Charlotte Soneson and Mark D. Robinson. Bias, Robustness And Scalability In Differential Expression Analysis Of Single-Cell RNA-Seq Data. bioRxiv, 2017a. URL http://biorxiv.org/content/early/2017/05/28/143289.

31. Davis J McCarthy, Yunshun Chen, and Gordon K Smyth. Differential expression analysis of multifactor RNA-Seq experiments with respect to biological variation. Nucleic acids research, 40(10):4288–97, may 2012a. ISSN 1362-4962. doi: 10.1093/nar/gks042. URL http://www.pubmedcentral.nih.gov/articlerender.fcgi?artid=3378882{&}tool=pmcentrez{&}rendertype=abstract.

32. Qiaolin Deng, Daniel Ramskold, Björn Reinius, and Rickard Sandberg. Single-cell RNA-seq reveals dynamic, random monoallelic gene expression in mammalian cells. Science (New York, N.Y.), 343(6167):193–6, jan 2014. ISSN 1095-9203. doi: 10.1126/science.1245316. URL http://www.ncbi.nlm.nih.gov/pubmed/24408435.

33. Daniel Bottomly, Nicole A R Walter, Jessica Ezzell Hunter, Priscila Darakjian, Sunita Kawane, Kari J Buck, Robert P Searles, Michael Mooney, Shannon K McWeeney, and Robert Hitzemann. Evaluating gene expression in C57BL/6J and DBA/2J mouse striatum using RNA-Seq and microarrays. PloS one, 6(3): e17820, jan 2011. ISSN 1932-6203. doi: 10.1371/journal.pone.0017820. URL http://www.pubmedcentral.nih.gov/articlerender.fcgi?artid=3063777{&}tool=pmcentrez{&}rendertype=abstract.

34. Xiaobei Zhou, Helen Lindsay, and Mark D Robinson. Robustly detecting differential expression in RNA sequencing data using observation weights. Nucleic acids research, 42(11):e91, jun 2014. ISSN 1362-4962. doi: 10.1093/nar/gku310. URL http://www.ncbi.nlm.nih.gOv/pubmed/24753412http://www.pubmedcentral.nih.gov/articlerender.fcgi?artid=PMC4066750.

35. Cameron Colin A. and Pravik K. Trivedi. No Title. In Regression analysis of Count Data, chapter 4.3. Cambridge University Press, Cambrdridge, second edition, 2013.

36. Johann a Gagnon-Bartsch and Terence P Speed. Using control genes to correct for unwanted variation in microarray data. Biostatistics, 13(3):539–52, 7 2012. ISSN 1468-4357. doi: 10.1093/biostatistics/kxr034. URL http://www.ncbi.nlm.nih.gov/pubmed/22101192.

37. Davide Risso, John Ngai, Terence P Speed, and Sandrine Dudoit. Normalization of RNA-seq data using factor analysis of control genes or samples. Nat Biotech, 32(9):896–902, 2014. ISSN 1087-0156. doi: 10.1038/nbt.2931. URL http://www.pubmedcentral.nih.gov/articlerender.fcgi?artid=4404308&tool=pmcentrez&rendertype=abstract.

38. Cole Trapnell, David G Hendrickson, Martin Sauvageau, Loyal Goff, John L Rinn, and Lior Pachter. Differential analysis of gene regulation at transcript resolution with RNA-seq. Nature biotechnology, 31(1): 46–53, jan 2013. ISSN 1546-1696. doi: 10.1038/nbt.2450. URL http://dx.doi.org/10.1038/nbt.2450.

39. Charlotte Soneson and Mark D Robinson. Towards unified quality verification of synthetic count data with countsimQC. Bioinformatics, oct 2017b. ISSN 1367-4803. doi: 10.1093/bioinformatics/ btx631. URL http://academic.oup.com/bioinformatics/article/doi/10.1093/bioinformatics/btx631/4345646/Towards-unified-quality-verification-of-synthetic.

40. Stephanie C Hicks, Mingxiang Teng, and Rafael A Irizarry. On the widespread and critical impact of systematic bias and batch effects in single-cell RNA-Seq data. bioRxiv, 2015. URL http://biorxiv.org/content/early/2015/12/27/025528.

41. Andrew Butler and Rahul Satija. Integrated analysis of single cell transcriptomic data across conditions, technologies, and species. doi.org, page 164889, jul 2017. doi: 10.1101/164889. URL https://www.biorxiv.org/content/early/2017/07/18/164889.

42. Alexey Sergushichev An algorithm for fast preranked gene set enrichment analysis using cumulative statistic calculation. bioRxiv, page 060012, jun 2016. doi: 10.1101/060012. URL https://www.biorxiv.org/content/early/2016/06/20/060012.

43. Dvir Aran, Zicheng Hu, and Atul J Butte. xCell: Digitally portraying the tissue cellular heterogeneity landscape. doi.org, page 114165, jun 2017. doi: 10.1101/114165. URL https://www.biorxiv.org/content/early/2017/06/15/114165.

44. Yaov Benjamini and Yosef Hochberg. Controlling the False Discovery Rate: A Practical and Powerful Approach to Multiple Testing. Journal of the Royal Statistical Society. Series B (Methodological), 57(1):289–300, 1995. URL https://www.jstor.org/stable/2346101?seq=1{#}page{_}scan{_}tab{_}contents.

45. Koen Van den Berge, Charlotte Soneson, Michael I. Love, Mark D. Robinson, and Lieven Clement. zingeR: unlocking RNA-seq tools for zero-inflation and single cell applications. bioRxiv, page 157982, jun 2017. doi: 10.1101/157982. URL https://www.biorxiv.org/content/early/2017/06/30/157982.

46. Dominic Grün, Lennart Kester, and Alexander van Oudenaarden. Validation of noise models for singlecell transcriptomics. Nature Methods, 11(6):637–640, apr 2014. ISSN 1548-7091. doi: 10.1038/nmeth. 2930. URL http://www.ncbi.nlm.nih.gov/pubmed/24747814http://www.nature.com/doifinder/10.1038/nmeth.2930.

47. Christoph Ziegenhain, Beate Vieth, Swati Parekh, Björn Reinius, Amy Guillaumet-Adkins, Martha Smets, Heinrich Leonhardt, Holger Heyn, Ines Hellmann, and Wolfgang Enard. Comparative Analysis of SingleCell RNA Sequencing Methods. Molecular Cell, 65(4):631–643, 2 2017. ISSN 10972765. doi: 10.1016/j. molcel.2017.01.023. URL http://linkinghub.elsevier.com/retrieve/pii/S1097276517300497.

48. Bhupinder Pal, Yunshun Chen, François Vaillant, Paul Jamieson, Lavinia Gordon, Anne C. Rios, Stephen Wilcox, Naiyang Fu, Kevin He Liu, Felicity C. Jackling, Melissa J. Davis, Geoffrey J. Lindeman, Gordon K. Smyth, and Jane E. Visvader. Construction of developmental lineage relationships in the mouse mammary gland by single-cell RNA profiling. Nature Communications, 8(1):1627, dec 2017. ISSN 2041-1723. doi: 10.1038/s41467-017-01560-x. URL http://www.nature.com/articles/s41467-017-01560-x.

49. Keisuke Fujita, Mitsuhiro Iwaki, and Toshio Yanagida. Transcriptional bursting is intrinsically caused by interplay between RNA polymerases on DNA. Nature Communications, 7:13788, dec 2016. ISSN 2041-1723. doi: 10.1038/ncomms13788. URL http://www.nature.com/doifinder/10.1038/ncomms13788.

50. Joseph N Paulson, O Colin Stine, Héctor Corrada Bravo, and Mihai Pop. Differential abundance analysis for microbial marker-gene surveys. Nature methods, 10(12):1200–2, sep 2013. ISSN 1548-7105. doi: 10.1038/nmeth.2658. URL http://www.ncbi.nlm.nih.gov/pubmed/24076764http://www.pubmedcentral.nih.gov/articlerender.fcgi?artid=PMC4010126http://www.nature.com/doifinder/10.1038/nmeth.2658http://dx.doi.org/10.1038/nmeth.2658.

51. Lizhen Xu, Andrew D Paterson, Williams Turpin, and Wei Xu. Assessment and Selection of Competing Models for Zero-Inflated Microbiome Data. PloS one, 10(7):e0129606, 2015. ISSN 1932-6203. doi: 10.1371/journal.pone.0129606. URL http://www.ncbi.nlm.nih.gov/pubmed/26148172http://www.pubmedcentral.nih.gov/articlerender.fcgi?artid=PMC4493133.

52. Davis J McCarthy, Yunshun Chen, and Gordon K Smyth. Differential expression analysis of multifactor RNA-Seq experiments with respect to biological variation. Nucleic acids research, 40(10):4288–97, may 2012b. ISSN 1362-4962. doi: 10.1093/nar/gks042. URL http://www.ncbi.nlm.nih.gov/pubmed/22287627http://www.pubmedcentral.nih.gov/articlerender.fcgi?artid=PMC3378882.

53. Catalina A. Vallejos, Davide Risso, Antonio Scialdone, Sandrine Dudoit, and John C. Marioni. Normalizing single-cell RNA sequencing data: challenges and opportunities. Nature Methods, page Under review, 2017.

54. Paul J. McMurdie and Susan Holmes. phyloseq: An R Package for Reproducible Interactive Analysis and Graphics of Microbiome Census Data. PLoS ONE, 8(4):e61217, apr 2013. doi: 10.1371/journal.pone. 0061217. URL http://dx.plos.org/10.1371/journal.pone.0061217.

55. Richard Bourgon, Robert Gentleman, and Wolfgang Huber. Independent filtering increases detection power for high-throughput experiments. Proceedings of the National Academy of Sciences ofthe United States of America, 107(21):9546–51, may 2010. ISSN 1091-6490. doi: 10.1073/pnas.0914005107. URL http://www.ncbi.nlm.nih.gov/pubmed/20460310http://www.pubmedcentral.nih.gov/articlerender.fcgi?artid=PMC2906865.

56. Charlotte Soneson and Mark D Robinson. iCOBRA: open, reproducible, standardized and live method benchmarking. Nature Methods, 13(4):283–283, mar 2016. ISSN 1548-7091. doi: 10.1038/nmeth.3805. URL http://www.nature.com/doifinder/10.1038/nmeth.3805.

57. Debarka Sengupta, Nirmala Arul Rayan, Michelle Lim, Bing Lim, and Shyam Prabhakar. Fast, scalable and accurate differential expression analysis for single cells. bioRxiv, 2016.

58. Mark A van de Wiel, Maarten Neerincx, Tineke E Buffart, Daoud Sie, and Henk MW Verheul. ShrinkBayes: a versatile R-package for analysis of count-based sequencing data in complex study designs. BMCBioinformatics, 15(1):116, 2014. ISSN 1471-2105. doi: 10.1186/1471-2105-15-116. URL http://bmcbioinformatics.biomedcentral.com/articles/10.1186/1471-2105-15-116.

59. Dirk F. Moore. Asymptotic Properties of Moment Estimators for Overdispersed Counts and Proportions. Biometrika, 73(3):583, dec 1986. ISSN 00063444. doi: 10.2307/2336522. URL http://www.jstor.org/stable/2336522?origin=crossref.

60. P. (Peter) McCullagh and John A. Nelder. Generalized linear models. Chapman and Hall, New York, second edition, 1989. ISBN 9780412317606. URL https://www.crcpress.com/Generalized-Linear-Models-Second-Edition/McCullagh-Nelder/p/book/9780412317606.

61. Simon N. Wood. Thin plate regression splines. Journal of the Royal Statistical Society: Series B (Statistical Methodology), 65(1):95–114, feb 2003. ISSN 1369-7412. doi: 10.1111/1467-9868.00374. URL http://doi.wiley.com/10.1111/1467-9868.00374.

62. A. Subramanian, P. Tamayo, V. K. Mootha, S. Mukherjee, B. L. Ebert, M. A. Gillette, A. Paulovich, S. L. Pomeroy, T. R. Golub, E. S. Lander, and J. P. Mesirov. Gene set enrichment analysis: A knowledge-based approach for interpreting genome-wide expression profiles. Proceedings of the National Academy of Sciences, 102(43):15545–15550, oct 2005. ISSN 0027-8424. doi: 10.1073/pnas.0506580102. URL http://www.pnas.org/cgi/doi/10.1073/pnas.0506580102.

